# Cell polarity linked to gravity sensing is generated by protein translocation from statoliths to the plasma membrane

**DOI:** 10.1101/2023.03.31.534658

**Authors:** Takeshi Nishimura, Shogo Mori, Hiromasa Shikata, Moritaka Nakamura, Yasuko Hashiguchi, Yoshinori Abe, Takuma Hagihara, Hiroshi Y. Yoshikawa, Masatsugu Toyota, Takumi Higaki, Miyo Terao Morita

**Author notes:** These authors contributed equally to this work (Takeshi Nishimura, Shogo Mori, Hiromasa Shikata).

## Abstract

Organisms have evolved under the gravitational force and sense the direction of gravity via statoliths in specialized cells. In the gravitropism of flowering plants, the starch-accumulating plastids, amyloplasts, in gravity sensing cells act as statoliths. The gravity sensing mechanism has long been considered a mechanosensing process by which amyloplasts transmit forces to intracellular structures, but the molecular support has not been reported. This study revealed that LAZY1-LIKE family proteins involved in gravity signaling in statocytes are localized to the amyloplast periphery and its proximal plasma membrane, resulting in polar localization according to the direction of gravity. We propose a gravity sensing mechanism by which LZY transmits the positional information of amyloplasts, i.e., the direction of gravity, by translocating to the plasma membrane.

## Main Text

Plants sense their inclination relative to the direction of gravity and reorient their growth direction accordingly, a process termed gravitropism (*1, 2*). The directional change is sensed mainly by specialized cells called statocytes, which contain plastids filled with dense starch granules termed amyloplasts (*3, 4*). The displacement of amyloplasts toward gravity as statoliths is believed to promote auxin transport in the direction of gravity, leading to organ bending (*5*). Several hypotheses have been proposed regarding the sensing mechanism by which the physical process of amyloplast displacement generates biochemical signals in statocytes, including the force sensing model (*6*) and position sensing hypothesis (*7*). However, the molecular mechanisms of gravity sensing and signaling remain largely unknown.

*LAZY1-LIKE* (*LZY*) family genes, which are conserved in most land plants, are involved in gravitropism and whole-plant architecture through the control of branch angles in various species in angiosperms, including trees (*8–15*). In Arabidopsis, LZY proteins have been shown to play critical roles in the signaling process controlling directional auxin flow following amyloplast displacement in statocytes in both shoots and roots (*16*). It should be noted that LZY3 is polarly localized to the plasma membrane (PM) in the direction of gravity and repolarized in response to the reorientation in statocytes, columella cells, in lateral roots (LRs) (*17*). This fact prompted us to investigate the mechanism of LZY3 polarization to understand gravity sensing and signaling in gravitropism.

First, we determined the regions responsible for membrane association in LZY3 protein because of the lack of detectable transmembrane domains. Basic–hydrophobic (BH) clusters in proteins can facilitate their membrane association in eukaryotes, including plants (*18–20*). A computational search indicated that all LZY proteins in Arabidopsis have more than two regions with a high BH score (≥ 0.6) that have the potential to serve as unstructured membrane-binding sites (fig. S1, A and B) (*18*). Gln substitutions were introduced at two sites, designated sites A and B, in LZY3 that decreased the BH scores of the corresponding sites (Fig. 1A, and fig. S1B). The expression of wild-type LZY3 fused with mCherry (LZY3-mCherry) and its mutants A-7Q, A-10Q, and B-7Q under the *LZY3* promoter rescued the primary root angle phenotypes of *lzy2;3;4*, which tended to grow upward, although their fluorescence signals were faint in the living columella cells of the primary roots (PRs; Fig. 1, B and C, and fig. S1, D and E). Both A-10Q and B-11Q exhibited significantly increased fluorescence intensity to different extents (fig. S1G). LZY3-mCherry carrying a mutation at site A localized to the PM and retained LZY3 activity, whereas that carrying a mutation at site B mainly localized in the cytosol and exhibited a variable growth direction (Fig. 1, C and D). Given that overexpression of LZY3-mCherry in statocytes resulted in PM localization without polarity and a random growth direction of LRs (*17*), B-11Q might have residual function. A-10Q;B-7Q exhibited PM localization with higher cytosolic fluorescence and a lower degree of rescue than A-10Q. The phenotypic difference between A-10Q and A-10Q;B-7Q might be consistent with the extent of polar localization (Fig. 1D, and fig. S1, C to F). The fluorescence signals of A10-Q;B-11Q were mostly found in the cytosol, and that expressing plants exhibited an almost random growth direction (Fig. 1, B and C). These suggest that sites B and A play major and minor roles in PM association, respectively, and that both regions help maintain LZY3 at low levels (fig. S1G). The existence of a putative PEST motif partly overlapping with site A implies its involvement in the regulation of LZY3 levels (fig. S2A) (*21, 22*). As expected, LZY3ΔPEST-mCherry lacking the PEST motif exhibited similar phenotypes of root growth and localization as A-10Q, implying quantitative regulation via proteasomal protein degradation (fig. S2, B and D). The addition of a farnesylation site [K8-far (*23*)] at the C-termini of B-11Q and A-10Q;B-11Q overrides the effect of their mutation on localization and activity, suggesting that the B-11Q and A-10Q;B-11Q mutants retain all functions of the wild-type LZY3 excluding association with the PM (fig. S3). In addition, the fact that the addition of K8-far reduced B-11Q levels implies a linkage between the membrane localization and quantitative regulation of LZY3.

**Fig. 1.**
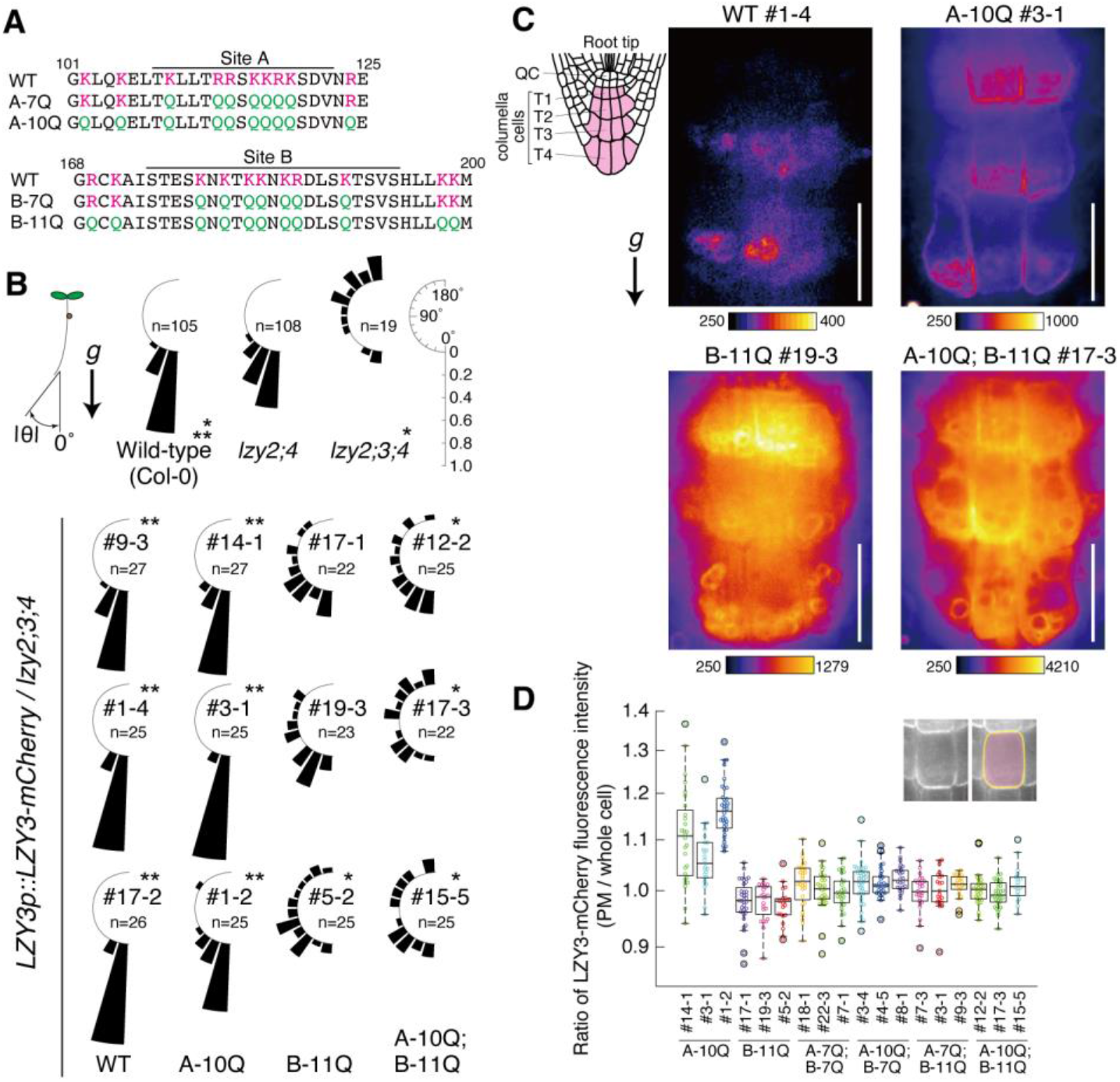
A potential membrane association site in LZY3 is crucial for subcellular localization in the columella cells and root gravitropism. (**A**) Potential plasma membrane association sites in LZY3. The lines indicate sites with basic– hydrophobic scores ≥ 0.6 according to computational prediction. In A-7Q and A-10Q, 7 and 10 basic amino acid residues at and near site A, respectively, were replaced with glutamine residues. In B-7Q and B-11Q, 7 and 11 basic residues at and nearby site B, respectively, were replaced with glutamine residues. The basic residues (K, R) and glutamine residue (Q) are colored magenta and green, respectively. (**B**) Growth angle of the primary root tips of 5-day-old seedlings of wild-type, *lzy2;4, lzy2;3;4*, and transgenic lines expressing mCherry-tagged LZY3 mutant proteins under the control of the *LZY3* promoter (*LZY3p::LZY3-mCherry*) in the *lzy2;3;4* mutants background. The direction of each root tip was measured as the absolute value toward the direction of gravity. The frequency was calculated as the proportion of root numbers that fell within intervals of 15° to the total number of analyzed roots for each line (range, 0–180°). Asterisks indicate significant differences [*P* < 0.0033 (0.05/15)] by Wilcoxon’s rank sum test with Bonferroni’s correction (*, compared to *lzy2;4;* **, compared to *lzy2;3;4*). (**C**) Representative confocal images of wild-type (WT), A-10Q, B-11Q, and A-10Q;B-11Q mutants of LZY3-mCherry in columella cells of primary roots. The area containing tier 1–3 columella cells of the representative lines is shown. The seedlings were kept vertically before and during imaging. The roots of the B-11Q and A-10Q;B-11Q lines were straightened at least 3 h before imaging to make the amyloplasts sediment toward the root tip, as those roots were meandering (fig. S1D). All images were acquired with an identical optical setting, and the color-coded heatmaps for signal intensities were placed below each image. Scale bars, 20 μm. *g*, the direction of gravity. (**D**) Ratio of fluorescence intensities of LZY3-mCherry at the plasma membrane (PM) to that of the whole cell. The ratios were calculated with mean values within a 3-pixel-width line drawn on the PM region and within the area surrounded by the line. For each line, 19–41 cells with clear cell edges in 5–8 roots were measured. WT, A-7Q, and B-7Q lines were not analyzed because of low protein accumulation.

Meanwhile, we noticed faint ring-shaped fluorescence signals in the tier 3 columella cells in wild-type LZY3-mCherry, which were observed at the position of amyloplasts. Given that LZY localizes to amyloplasts, LZY might help link gravity sensing to the signaling. However, the weak fluorescence intensity of LZY3-mCherry made it difficult to analyze the behavior of LZY3 in tier 2 cells, which are the major contributors to gravitropism (*24*). Then, we examined whether LZY4, which redundantly functions in root gravitropism (*25, 26*), can be applied for live cell imaging analysis. There are four splicing variants of LZY4 (annotated in TAIR10), three of which lack the CCL region (fig. S4, A and B), which is important for LZY1/2/3 function (*16, 17, 27*). We demonstrated that LZY4.4 with a CCL-like sequence is the functional form in both gravitropism and interactions with the BRX domain of RLDs in the Y2H system (fig. S4, C and D). RLD1 is an interacting partner of LZY in the regulation of auxin transport via membrane trafficking, and it is recruited to the PM in a LZY2/3-dependent manner (*17, 28*). LZY4.4 recruited RLD1 on the PM in Arabidopsis protoplasts (fig. S5). In addition, *LZY4.4-mClover3* driven by the statocyte-specific *ADF9* promoter rescued the PR angle phenotype of *lzy1;2;3;4* (fig. S6). These results indicate that the molecular function of LZY4.4 (hereafter LZY4) is almost equivalent to that of LZY3 in root statocytes.

Ring-shaped and linear fluorescence signals from functional *LZY4p::LZY4-mScarlet* was observed in the tier 2 columella cells of young LRs (Fig. 2, A and B). Because LZY4-mCherry localized to the PM in protoplasts (fig. S5B), the linear fluorescence signal of LZY4-mScarlet along cell contours is likely to occur on the PM of statocytes. Notably, LZY4-mScarlet on the PM was observed only proximal to amyloplasts, exhibiting apparent polarity to the direction of gravity (Fig. 2A). LZY4 also has a basic amino acid-rich region in the middle of the protein (BH score ≥ 0.6; fig. S7A). LZY4-mScarlet carrying the B-6Q mutation in this region exhibited lower PM localization and weaker capacity for rescue, suggesting a PM-binding mechanism similar to LZY3 and its functional importance (Fig. 2, B and C, and fig. S7B). The ring-shaped signal merged with the plastid marker PIC1-GFP, indicating that LZY4 localized to the amyloplast periphery (Fig. 2D). LZY4-mScarlet lacking the N-terminal domain I (LZY4-ΔI-mScarlet) was present at the PM and in the cytosol, but the ring-shaped signal was hardly detected (Fig. 2E). In addition, LZY4-ΔI-mScarlet was not fully functional (Fig. 2B). Consistently, mCherry fused with the N-terminal 54 amino acids of LZY3, including the conserved domain I, displays obvious ring-shaped signals (fig. S8). These findings suggest that the N-terminal region including domain I is involved in the amyloplast localization of LZY3/4. It has been reported that several mutations in genes encoding components of the translocon at the outer chloroplast membrane (TOC) complex enhance the phenotype of *altered response to gravity 1* (*arg1*) and its paralog *arg1-like2* (*arl2*) (*29, 30*). *ARG1* and *ARL2*, encoding J-domain proteins, have been revealed to be involved in gravity signaling in statocytes (*31–33*) similarly as *LZYs*. We investigated the possible involvement of ARG1/ARL2 in the plastid localization of LZY4-mScarlet. In the *arg1* and *arl2* background, the LZY4-mScarlet signal became undetectable in amyloplasts and at the PM, and the signal was dispersed in the cytosol (Fig. 2F, and fig. S9). ARG1 and ARL2 are likely to be involved in the recruitment of LZYs to amyloplasts and the PM, possibly through their chaperone activity. Considering that the precursor proteins targeted to plastids interact with TOC via their N-terminal transit peptides with the help of molecular chaperones (*34*), mechanisms similar to plastid protein targeting could be involved in targeting LZYs to amyloplasts.

**Fig. 2.**
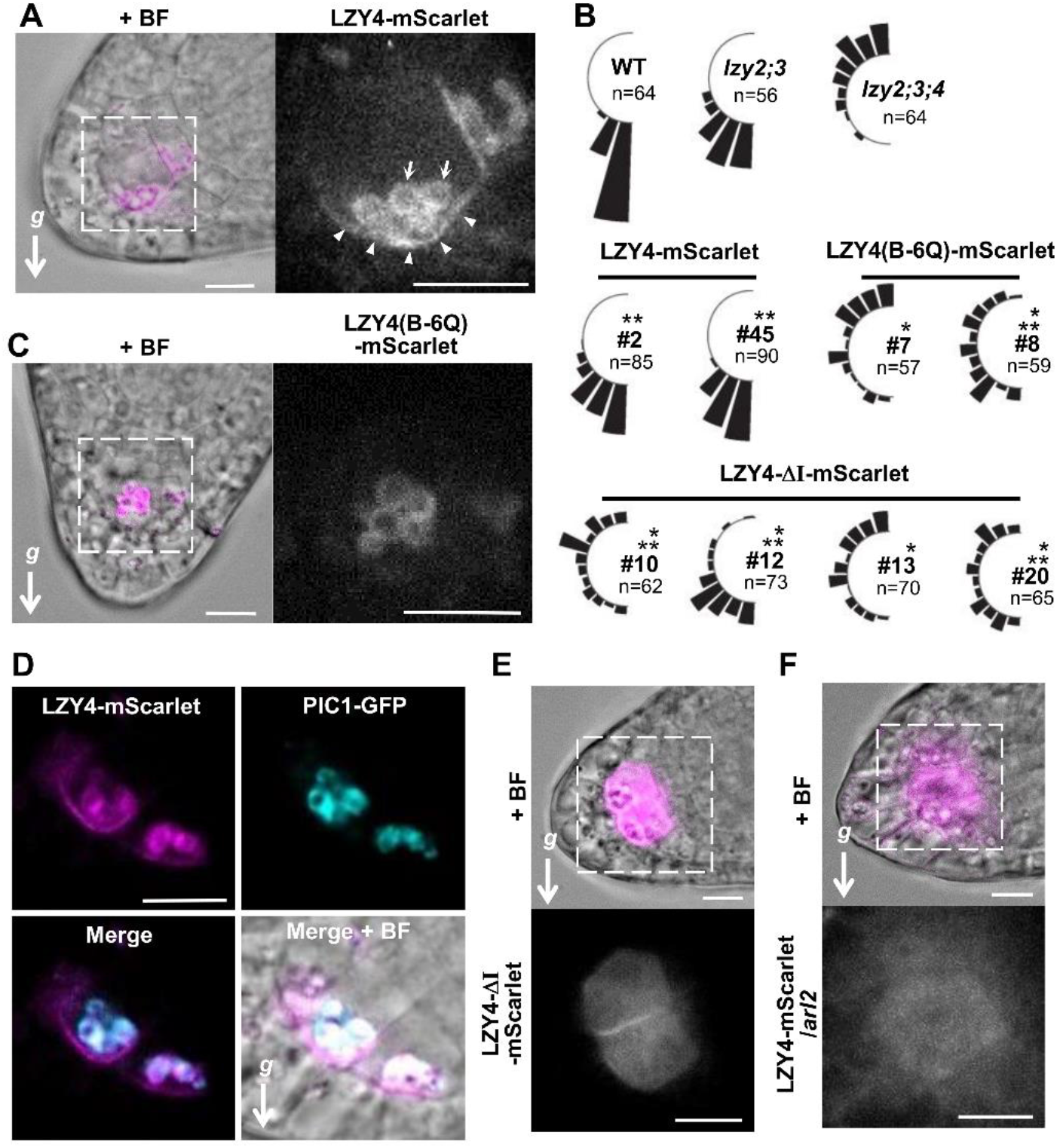
Amyloplast and basal plasma membrane localization of LZY4 in the columella cells. **(A)** Subcellular localization of LZY4-mScarlet (#2) / *lzy4* in lateral root columella cells. Arrows indicate the amyloplast-like ring shape localization of LZY4-mScarlet. Arrowheads indicate localization to the basal plasma membrane. Scale bars, 10 μm. **(B)** Complementation test of *LZY4-mScarlet* / *lzy2;3;4, LZY4(B-6Q)-mScarlet* / *lzy2;3;4*, and *LZY4-ΔI-mScarlet* / *lzy2;3;4* in the direction of the primary root tips of 5-day-old seedlings. Asterisks indicate significant differences [*P* < 0.0045 (0.05/11)] by Wilcoxon’s rank sum test with Bonferroni’s correction (*, compared with *lzy2;3;* **, compared with *lzy2;3;4*). **(C)** Subcellular localization of LZY4(B-6Q)-mScarlet (#7) / *lzy2;3;4* in lateral root columella cells. Scale bars, 10 μm. **(D)** Co-localization of LZY4-mScarlet and PIC1-GFP in amyloplasts. Scale bar, 10 μm. **(E)** Subcellular localization of LZY4-ΔI-mScarlet (#12) / *lzy2;3;4*. Scale bars, 10 μm. **(F)** Subcellular localization of LZY4-mScarlet (#2) in the *arl2* mutant background. Scale bars, 10 μm. *g*, the direction of gravity.

Next, we examined the effect of gravistimulation by reorientation on the localization of LZY4-mScarlet through live cell imaging of LR columella cells using a vertical stage confocal microscope (*35*). Before gravistimulation, polar localization of LZY4-mScarlet on the PM was maintained throughout the observation, although amyloplasts were slightly agitated (fig. S10B, and movie S1). After gravistimulation, LZY4-mScarlet appeared on the PM of the new bottom side of the cells at almost the same time as amyloplast displacement, and the signal gradually became stronger (Fig. 3, A and B, fig. S10C, and movie S2). Simultaneously, the LZY4-mScarlet signal on the PM in the original direction of gravity gradually disappeared, resulting in the generation of the new polarity of LZY4. The starchless *pgm* mutant has amyloplasts that hardly sediment because of the lack of dense starch granules, and it exhibits reduced gravitropism (*36–38*). LZY4-mScarlet localized to smaller starchless amyloplasts and the PM in the *pgm* mutant. Notably, LZY4-mScarlet failed to accumulate polarly on the PM in the *pgm* mutant, probably because of the movement of amyloplasts (Fig. 3C, and movie S3), indicating that amyloplast sedimentation is required for the polarization of LZY4 on the PM.

**Fig. 3.**
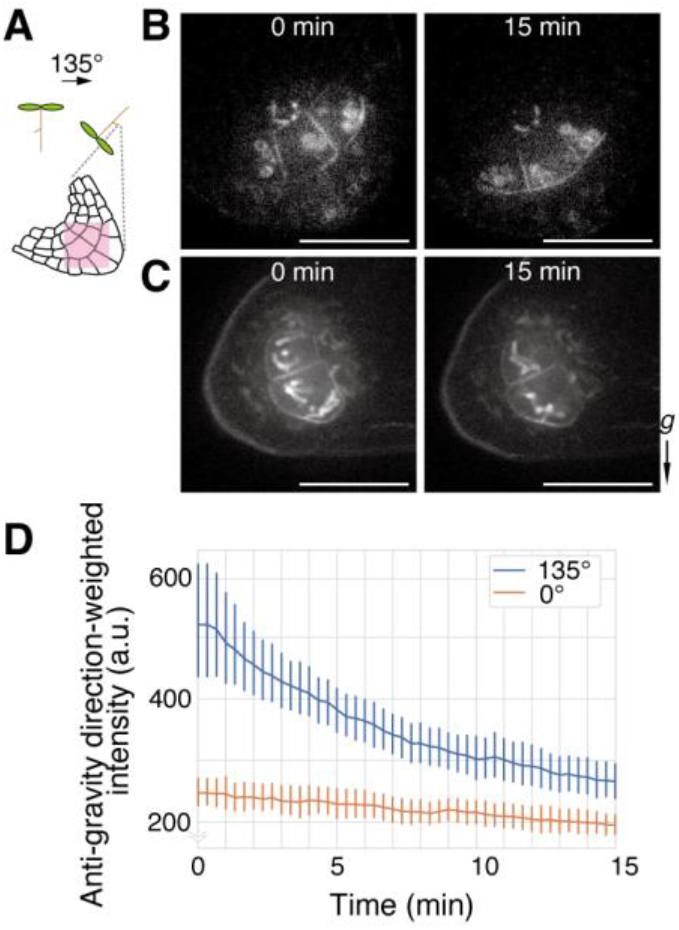
Polar localization of LZY4 changes in response to gravistimulation. **(A)** Gravistimulation and the site of observation. Stage 2 lateral roots (< 3 mm) were used. **(B** and **C)** Representative confocal images of the columella cells of *LZY4p::LZY4-mScarlet* in the *lzy4* **(B)** or *lzy4 pgm-1* **(C)** background immediately before (0 min) or 15 min after gravistimulation. The seedlings were maintained in a vertical position before and during imaging. mScarlet was excited at 561 nm, and the fluorescence that passed through a 617/73 nm emission filter was detected. *g*, the direction of gravity. Scale bars, 25 μm. **(D)** Temporal changes in the localization of LZY4-mScarlet on the plasma membrane along the direction of gravity. Images were taken at 20-s intervals for 15 min. n = 8 (0°) and n = 17 (135°). Error bars, standard deviation.

The temporal change in the localization of the LZY4-mScarlet signal on the PM along the direction of gravity was quantified (Fig. 3D, and fig. S11). Briefly, the fluorescence intensity along the vertical line (y-axis) is integrated along the x-axis and weighted by the opposite direction of gravity. Thus, a higher value of weighted intensity means that LZY4-mScarlet polarizes at a higher position, whereas a lower value means that LZY4-mScarlet polarizes in the direction of gravity. Without gravistimulation, the weighted intensity gradually and slightly decreased, probably because of quenching during the 15-min observation. By contrast, the value decreased significantly after gravistimulation, indicating repolarization of LZY4-mScarlet in the direction of gravity. An incremental decrease in the value occurred immediately after gravistimulation, presumably linked with amyloplast displacement. To clarify the temporal relationship between the repolarization of LZY4 and other processes in gravitropism, we examined each event with primary roots under the same conditions as much as possible (fig. S12). The behavior of LZY4-mScarlet in PRs was nearly the same as that in LRs, excluding the lower fluorescence intensity. Significant repolarization of LZY4-mScarlet was detected within 15 min after gravistimulation (fig. S12, A to C, and movie S4). The asymmetric distribution of auxin detected using the ratiometric auxin biosensor R2D2 (*39*) appeared 16 min after gravistimulation, and it became significant at 30 min (fig. S12, E and F). Significant curvature appeared 30 min after gravistimulation (fig. S12D). In addition, amyloplast displacement occurred 3 min after gravistimulation in our observation system, as previously reported (*16*). Therefore, repolarization of LZY4-mScarlet can be described as a chronologically crucial process linking gravity sensing to signaling. As reported previously, the polar pattern of PIN3-GFP in the root cap columella after gravistimulation was observed much later in our system (*17*). This implies that the LZY-RLD module may not directly regulate the trafficking of PIN3 but rather other regulatory factors for auxin transport.

Given that LZY interacts electrostatically with negatively charged lipids, the lipids might polarly distribute on the PM and contribute to the polar localization of LZYs. Therefore, we examined the distribution of several negatively charged lipids in the statocytes of PRs and LRs using fluorescent biosensors (fig. S13). Only the phosphatidylinositol-4,5-bisphosphate biosensor CITRIN-2xPH^PLC^ (*40*) tended to be polarly distributed toward the tip. However, clear repolarization of CITRIN-2xPH^PLC^ was not detected upon gravistimulation in the examined time window. The distribution of these lipids is unlikely to be related to the polarity of LZYs. Next, to elucidate the relationship between LZY4 residing on amyloplasts and its polarization on the PM, the behavior of LZY4 was analyzed using the *LZY4p::LZY4-mEos2/lzy4* rescued line (fig. S14A). LZY4-mEos2 on the amyloplasts was photoconverted from green to red by irradiation with a 405-nm laser on the vertical stage confocal microscope. The red fluorescence signal was observed only in amyloplasts immediately after irradiation, and linear red fluorescence emerged on nearby PMs (Fig. 4A, fig. S14B, and movie S5). The fluorescence intensity on the PM proximal to amyloplasts significantly increased 1 min after irradiation (Fig. 4B). Although we could not dismiss the possibility that a trace amount of cytoplasmic LZY4-mEos2 was photoconverted and accumulated at the PM, this result suggests that LZY4 rapidly translocates from amyloplasts to the PM. Then, it is expected that the position of amyloplasts determines the LZY4 accumulation site on the PM independent of the direction of gravity. To test this possibility, amyloplasts were manipulated by an optical tweezer during observation by confocal laser scanning microscopy (*41*). After trapping some amyloplasts, the amyloplasts were manipulated to put them in close proximity to the PM (figs. S15 and S16, and movies S6 and S7). The fluorescence intensity of LZY4-mScarlet significantly increased at the PM region where the amyloplasts were newly placed nearby (Fig. 4, C and D, and movie S8), indicating that amyloplasts act as determinants of LZY4 polarity on the PM.

**Fig. 4.**
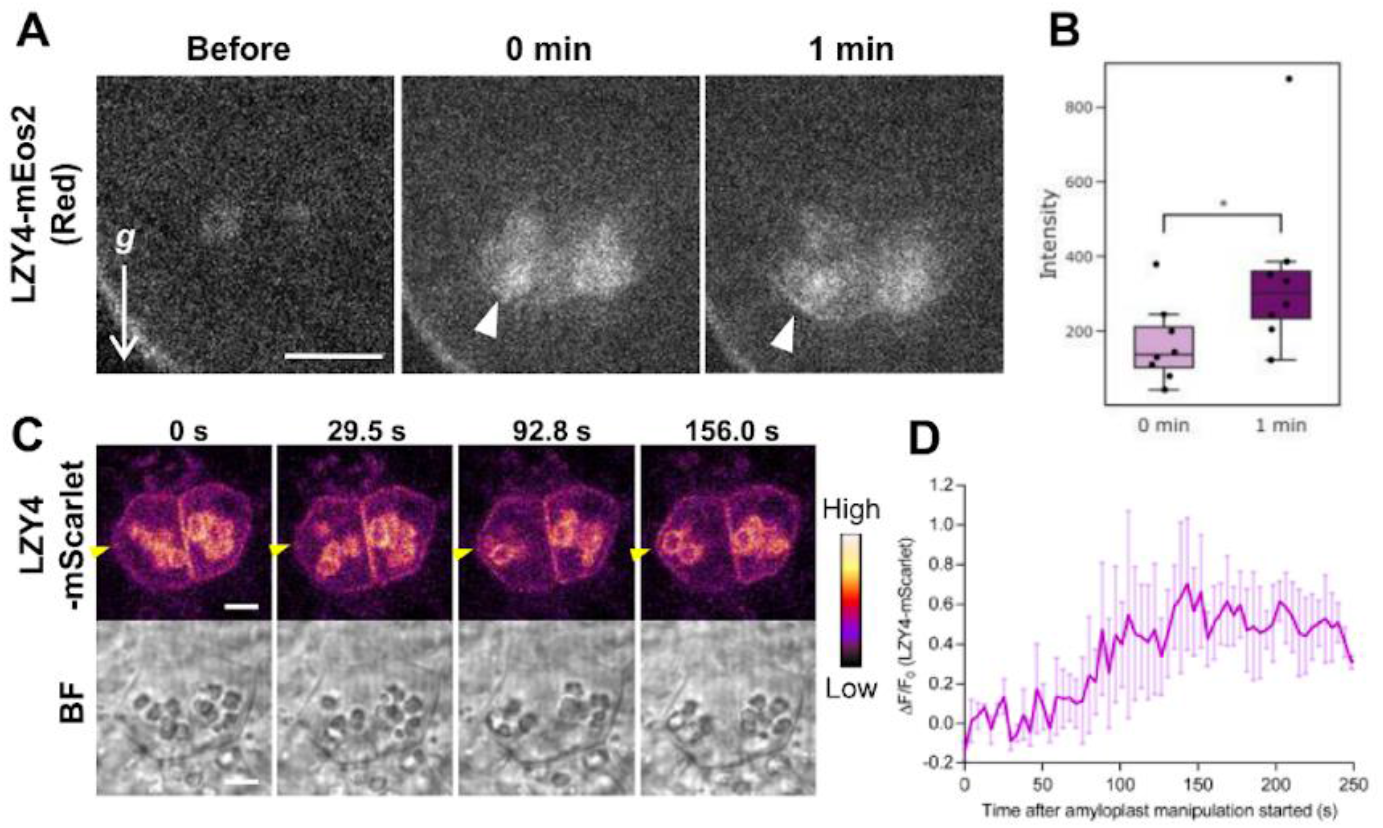
Translocation of LZY4 from the amyloplast to the plasma membrane (PM). **(A** and **B)** Observation of the translocation of LZY4-mEos2 from the amyloplasts to the PM by the photoconversion method **(A)**. Immediately after photoconversion, there was almost no red fluorescence signal on the PM (white arrowhead in Red; 0 min). At 1 min after photoconversion, a linear red fluorescence signal was observed at the PM in close proximity to the amyloplasts (white arrowhead in Red; 1 min). Scale bar, 10 μm. *g*, the direction of gravity. There was a significant difference between the findings at 0 and 1 min (Wilcoxon’s rank sum test, *P* < 0.05). The increase in the red fluorescence signal on the PM at 1 min after photoconversion was quantitatively analyzed **(B)**. n = 8. See also fig. S14B showing images of mEos2 green fluorescence signals. **(C** and **D)** Temporal changes in the localization of LZY4-mScarlet upon manipulating amyloplasts with an optical tweezer. Amyloplasts were fixed with the optical tweezer and positioned near the PM (yellow arrowheads) by moving the roots immobilized on the stage. The representative change is shown in **C.** Scale bar, 5 μm. The temporal changes in the signal of mScarlet on the PM where the amyloplasts were positioned are quantitatively shown in **D**. n = 3.

Gravity sensing in gravitropism has long been considered a mechanosensing mechanism whereby the weight of statoliths exerts a force on intracellular structures (*6, 42*). Based on the experimental results using angiosperm shoots, it has recently been suggested that the sensor functions as a clinometer but not as a force sensor (*43*). This supports the ‘position sensor hypothesis,’ which proposes that proximity or contact between statoliths and the membrane induces local auxin fluxes (*7*) and that statocytes act as clinometers (*44*), although there are no known molecules to support this hypothesis. We revealed in this study that LZY exhibits behavior that fits the position sensor hypothesis well. LZY polarity on the PM was formed according to the position of amyloplasts by translocation of LZY from amyloplasts to the PM. LZY appears to act as a signal molecule that transmits the positional information of statoliths to the PM, which directly links gravity sensing to subsequent signaling processes. Polarly located LZY recruits RLD to promote polar auxin flow in the direction of gravity (*17*).

## Acknowledgments

We are grateful to Dr. Nozomi Kawamoto (National Institute for Basic Biology) and Dr. Masahiko Furutani (Kumamoto University) for helpful discussions and to Dr. Tomohiro Uemura (Ochanomizu University) for providing Deep cell. *arg1-3* was a kind gift from Dr. Patrick Masson (University of Wisconsin-Madison). mRFP1-Spo20p-PABD was a kind gift from Dr. Martin Potocký (Czech Academy of Sciences). The support of plant cultivation rooms was provided by the Model Plant Research Facility of National Institute for Basic Biology. We also thank Wakana Takase, Hiroe Motomura, Kikumi Miyoshi, Mayako Hamada, Yuka Yamada, and Yoriko Soma for technical assistance; the Salk Institute Genomic Analysis Laboratory for providing the sequence-intexed Arabidopsis T-DNA insertion mutants; and the Arabidopsis Biological Resource Center and GABI-Kat for providing seeds of the *A. thaliana* T-DNA insertion mutants and lipid biosensor lines (P5R and P24Y).

## Funding

Japan Society for the Promotion of Science (JSPS) KAKENHI, Grant-in-Aid for Scientific Research 19H03254 (MTM)

Ministry of Education, Culture, Sports, Science and Technology (MEXT) KAKENHI, Grant-in-Aid for Scientific Research on Innovative Areas 18H05488 (MTM)

Japan Science and Technology Agency (JST), Core Research for Evolutionary Science and Technology (CREST) JPMJCR14M5 (MTM)

Takeda Science Foundation (MTM)

Naito Foundation (MTM)

JSPS KAKENHI, Grant-in-Aid for Scientific Research 22H00302 (HYY)

JSPS KAKENHI, Grant-in-Aid for Challenging Exploratory Research 20K21117 (HYY)

JSPS KAKENHI, Grant-in-Aid for Scientific Research on Innovative Areas 18H05492 (TH)

JSPS KAKENHI, Grant-in-Aid for Scientific Research 20H03289 (TH)

JST, CREST JPMJCR2121 (TH)

JSPS KAKENHI, Grant-in-Aid for Early-Career Scientist 18K14731 (MN)

JSPS KAKENHI, Grant-in-Aid for Early-Career Scientist 20K15826 (HS)

## Author contributions

Conceptualization: MTM, TN, HS

Design of Experiments: MTM, TN, HS, MT

Methodology: HYY, MT

Investigation: TN, HS, SM, MN, YA, TH, YH, TH

Funding acquisition: MTM, MN, HS

Project administration: MTM

Supervision: MTM

Writing: MTM, TN, HS, MT, TH

## Supplementary Materials

Materials and Methods

Figs. S1 to S16

References (*45–60*)

Movies S1 to S8

Table S1

