## Supplementary material for "Cell polarity linked to gravity sensing is generated by protein translocation from statoliths to the plasma membrane": sup.figs

### Supplementary Materials

#### Materials and Methods

Plant materials and growth condition

*Arabidopsis thaliana* (L.) Heynh. accession Columbia-0 (Col-0) was used as the wild-type. The following mutant alleles and marker lines were used: *lzy2*, *lzy3* (*16*), R2D2 (*39*), *pgm-1* (*45*), *arg1-3*, *arl2-4* (*33*), P5R and P24Y (*40*). *lzy4-3* (GABI_479C08) was obtained from Nottingham Arabidopsis Stock Centre (NASC) and backcrossed with Col-0 four times prior to the construction of multiple mutants, which is designated as *lzy4* here. *LZY4p::LZY4-mScarlet* / *lzy4* was crossed with mutants to introduce the transgene. Surface-sterilized seeds were sown on MS plates [1 x Murashige Skoog salts, 1% (w/v) sucrose, 0,01% myoinositol, 0.05% (w/v) MES (2-(N-morpholino) ethanesulfonic acid), and 0.5% (w/v) gellan gum; pH 5.8], incubated in the dark at 4°C for 3 days, and then grown vertically at 23°C in a growth chamber under continuous light condition (80 ~ 120 µmol/$m^2$s) for 10 to 14 days, and replanted on the soil as necessary (*17*).

Plasmid construction

To efficiently get transformants in *lzy2;3;4* triple mutants background, the pGWB501(*LZY3p::LZY3-mCherrry*) construct (*46*, *16*) was modified. Two *NOS* terminator fragments downstream of *LZY3-mCherry* or Hyg-resistance gene *hpt* were replaced with native *LZY3* terminator or rbcS-E9 terminator from pTA7002 vector (*47*), respectively. Those fragments were PCR-amplified with primers (#1-6) and assembled with In-Fusion HD Cloning Kit (Clontech). Besides, first two exons of *LZY3* gene were introduced into the coding region of *LZY3* cDNA by inserting a *Xho*I/*Nru*I-digested fragment from pFAST-R01 (*LZY3g*) (*16*). To introduce mutations at the site A and B of *LZY3* on the *LZY3p::LZY3-mCherry* vector, a fragment containing respective mutation was produced with PCR amplification using primers (#7-9 for A-7Q, #9-11 for A-10Q, #12-14 for B-7Q, or #14-16 for B-11Q) and then replaced with the wild-type sequence of *LZY3* in pUC19(*35S::LZY3-mCherry*) (*17*) by assembling with a PCR fragment using primers (#17 and #18 for site A, or #18 and #19 for site B). Finally, those mutated *LZY3-mCherry* were introduced into *Nru*I/*Sac*I sites of the modified pGWB501 construct. To delete the potential PEST motif on LZY3, PCR amplification using primers (#20 and #21) was performed with the modified pGWB501(*LZY3p::LZY3-mCherrry*) as a template, followed by self-ligation. The PEST-deleted *LZY3-mCherry* was re-introduced into *Nru*I/*Sac*I sites of the modified pGWB501 construct. To introduce the farnesylation motif (SKDGKKKKKKSKTKCVIM) from K-Ras (*48*) at the C-terminus of LZY3-mCherry, PCR amplification was performed using primers (#22 and #24 for $1^{st}$ PCR, #24 and #25 for $2^{nd}$ PCR) and pUC19(*35S::LZY3-mCherry*) including mutations as templates. The resulting *LZY3-mCherry-K8-far* was introduced into *Nru*I/*Sac*I sites of the modified pGWB501 construct. The constructs were introduced using floral dip into *lzy2;3;4* triple mutants (*49*).

The YFP- and mCherry-coding sequences were amplified with primers #26-28 and primers #29 and #30, respectively, and the products that have a mutation at the *Aau*I site on their coding sequences were digested with *EcoR*I and *Aau*I. In addition to each fragment, an *Aau*I/*Not*I-digested fragment including the *CaMV 35S* terminator (*35St*) from the pExtag vector (*50*) was inserted into the pGreenII0229 vector. The resulting vectors were designated as pGreenII0229_*YFP-(AauI)::35St* and *mCherry-(AauI)::35St*, respectively. The coding sequences of C2 domain from bovine Lactadherin (274-431 a.a., LactC2) (*51*) and KA1 domain from yeast MARK1 (901-1037 a.a., KA1$^{MARK1}$) (*52*) was gene-synthesized and inserted into *Aau*I site of pGreenII0229_*mCherry::35St* and *YFP-(AauI)::35St* vectors, respectively. The fragment including a coding sequence of the farnesylation motif was produced with primers (#31 and #32) and then inserted into pGreenII0229_*mCherry-(AauI)::35St*. The PA-biosensor mRFP1-Spo20p-PABD (*53*) was amplified by PCR with primers (#33 and #34) and then inserted into the *Eco*RI and *Xba*I-sites of the pGreenII0229_*35St* vector. The open reading frame of *LTI6b* was PCR amplified with using primers (#35 and #36 for 1$^{st}$ PCR, #37 and #38 for 2$^{nd}$ PCR) and genomic Arabidopsis DNA as a template, and then assembled with *Aau*I-digested pGreenII0229_*YFP-(AauI)::35St* with the In-Fusion kit. The coding sequence of N-terminal region of LZY3 (1-54 a.a.) was PCR amplified using primers (#39 and #40) and assembled with *Eco*RI-digested pGreenII0229_*mCherry-(AauI)::35St* with NEBuilder HiFi DNA Assembly kit (NEB). Stop codon was introduced into the *Aau*I site using annealed oligos (primers #41 and #42). The *Eco*RI/*Hind*III-digested fragment of *ADF9* promoter from pFAST-R01 (*ADF9p::LZY3)* was first inserted into *the* pGreenII0229_*35St* vector and then digested with *Eco*RI. The digested fragment including *ADF9* promoter was introduced into pGreenII0229(*mCherry-LactC2*, *YFP-KA1$^{MARK1}$*, *mCherry-K8-far*, *mRFP1-Spo20p-PABD*, *YFP-LTI6b*, and *LZY3(1-54)-mCherry*). The constructs were introduced using floral dip into Col-0 plants (*49*). Unless specified otherwise in the figure legend, T3 generations of the produced lines were used for the described experiments.

We used the Gateway Cloning System (Invitrogen) to construct *gLZY4-1*, *gLZY4-2, gLZY4-3*, *gLZY4-4*, *LZY4p::LZY4-mScarlet*, *LZY4p::LZY4(B-6Q)-mScarlet*, *LZY4p::LZY4ΔI-mScarlet*, *LZY4p::LZY4-mEos2*, *ADF9p::LZY4-mClover3*, and *ADF9p::PIC1-GFP*. pmmScarlet_C1 was a gift from Dorus Gadella [Addgene plasmid # 85042(*54*)]. pNCS-mClover3 was a gift from Michael Lin [Addgene plasmid # 74236(*55*)]. pRSETa mEos2 was a gift from Loren Looger [Addgene plasmid # 20341(*56*)]. The genomic regions specifically expressing *LZY4.1* (#43 and #44), *LZY4.2* (#43 and #45), *LZY4.3* (#43 and #46), and *LZY4.4* (#43 and #47) from 584 bp upstream from the start codon of *LZY4* were amplified by PCR and introduced into the pENTR vector containing *NOS* terminator by ligation with the restriction sites *Bam*HI and *Sac*I. Those vectors were then introduced into the pGWB501 binary vector (*46*) using LR clonase (Thermo Fisher Scientific). *LZY4p::LZY4-mScarlet* was created by amplifying a 584-bp upstream fragment from the start codon of *LZY4* (#48 and #49) and the full-length cDNA of *LZY4.4* (#50 and #51) by PCR and introducing them into pENTR-*mScarlet* by In-fusion HD Cloning Kit (In-Fusion, Clontech). *LZY4ΔI-mScarlet* was produced by inverse-PCR using *pENTR-LZY4p::LZY4-mScarlet* as a template to delete 36 bp after the start codon of *LZY4* (#52 and #53) and ligation it with restriction site *Spe*I. For *LZY4(B-6Q)-mScarlet*, a 96-bp (229-324 bp) fragment of the CDS of *LZY4*, in which the adenine at 247, 250, 253, 274, 277, and 304 was replaced to cytosine, was amplified using primers introducing the mutation by PCR (#54, #55, #56, #57 and #58). The PCR fragment was then replaced with wild-type CDS on pENTR-*LZY4p::LZY4-mScarlet* using NEBuilder HiFi DNA Assembly. For *LZY4p::LZY4-mEos2*, the CDS of *mEos2* were amplified by PCR (#59 and #60) and replaced with mScarlet on *LZY4p::LZY4-mScarlet* using In-Fusion. The CDS of *PIC1* (*57*) was amplified by PCR (#61 and #62) and introduced into pENTR-*ADF9p::GFP* by ligation between the *ADF9* promoter and GFP with the restriction site *Sph*I. The CDS of *LZY4* was also amplified by PCR (#63 and #64) and introduced between the *ADF9* promoter and *mClover3* of pENTR-*ADF9p::mClover3* by In-Fusion. These vectors were then introduced into the pGWB501 binary vector using LR clonase. The constructs of *gLZY4-1*, *gLZY4-2*, *gLZY4-3*, *gLZY4-4*, *LZY4p::LZY4-mEos2*, and *ADF9p::LZY4-mClover3* were introduced into *lzy1;2;3;4* mutant, and *LZY4p::LZY4-mScarlet*, *LZY4p::LZY4(B-6Q)-mScarlet,* and *LZY4p::LZY4ΔI-*

*mScarlet* were introduced into *lzy2;3;4* mutant, and *ADF9p::PIC1-GFP* was introduced into Col-0.

For transient assay using Arabidopsis protoplasts, the CDS of *LZY4* (#65 and #66) and the 3'-terminal 258-bp fragment of *RLD1* CDS, which is corresponding to the BRX-domain were amplified by PCR and their fragments were fused with GFP/mCherry genes containing *NOS* terminator on the pUC19 vector under the *35S* promoter (*35S::LZY4.4-mCherry* and *35S::RLD1BRXd-GFP*). *35S::GFP*, *35S::mCherry*, *35S::mCherry/GFP-LTI6b*, and *35S::RLD1-GFP* were described previously (*17*).

For interaction assay using Y2H, the region corresponding to CCL-Like in LZY4.4 (487-531 bp) was amplified by PCR (#67 and #68) and introduced into the MCS of pGADT7 by In-fusion. RLD1-C (2,077-3,312 bp) (#69 and #70), RLD1-CΔBRX (2,077-3,057 bp) (#69 and #71), and RLD1-BRX (3,052-3,240 bp) (#70 and #72) were amplified by PCR and introduced by In-fusion into the MCS of pGBKT7.

Quantification of primary root growth angle

For analyses of the gravitropic phenotype of primary roots in wild-type, *lzy* mutants, and transgenic plants, the seedlings were grown on the MS plates kept vertically for 5 days. Photographs were taken with a general photo scanner, and the angle between the direction of gravity and root tip growth was measured using the ImageJ software (https://imagej.nih.gov/ij/). For statistical analyses, we assessed the quality of variances for the data by Levene's test and found differences between the variances in most cases. Hence, the significant differences among genotypes were evaluated by the Kruskal-Wallis rank sum test followed by the Wilcoxon rank sum test with Bonferroni's correction ($P < 0.05/N$; $N$, number of genotypes).

Live-imaging of fluorescent proteins in columella cells

To image fluorescent proteins in columella cells of primary or lateral roots, 5-day-old or 7-day-old seedlings, respectively, grown on the vertically-kept plates were quickly transferred into a water droplet on the imaging glass slide and were carefully covered with a cover slip. The imaging slides were mounted on the 360˚-rotatable vertical stage of the confocal microscope equipped with the spinning disk confocal scanning unit and the back-illuminated EM-CCD camera (*58, 35*). A 60x silicone immersion objective lens (UPLSAPO60XS2, 1.3 N.A., Olympus) was used for all image acquisitions. YFP, CITRIN, and steady-state mEos2 (green fluorescence) were excited at 488 nm and the fluorescence passed through a 525/50 nm emission filter was detected. mCherry, mScarlet, mRFP1, and photoconverted mEos2 (red fluorescence) were excited at 561 nm and the fluorescence passed through a 600/37 nm emission filter was detected, unless specified otherwise in the figure legend. Note that lateral roots at stage 2 (*17*) were observed.

To generate time-lapse movies, the positions of samples on images acquired at each time point were adjusted with the "Auto Align" algorithm of the MetaMorph software (Molecular Device) by referring to the bright field images acquired at the one previous time point.

Interaction assay using Y2H

MatchmakerTM Gold Yeast Two-Hybrid System Kit (Clontech) was used for interaction assay. The CCL-Like region for LZY4.4 was fused to the C-terminus of GAL4 activation domain (AD) of pGADT7, and then the plasmid was transformed into Y137 yeast strain (Clontech) using FastTM-Yeast Transformation Kit (G-Biosciences). The truncated RLD1 was fused to the C-terminus of GAL4 DNA-binding domain (DBD) of pGBKT7 and then the plasmids were transformed into Y2HGold yeast strain (Clontech). After mating, spot assays were performed on -Leu/-Trp/-His/-Ade/+Aureobasidin A (125 ng/ml) (TaKaRa) media by incubating for 3 days at 30°C.

Transient assay

Transient assays with protoplasts of Arabidopsis suspension culture were carried out (*17*). Plasmids were introduced into protoplasts, which were generated from suspension culture cells, containing 0.4 M mannitol and 32% (w/v) PEG6000 (Nacalai tesque). After 12-18 h incubation at 23°C, confocal images of GFP and mCherry fluorescence were obtained with a confocal laser scanning microscopy (FV-1000; Olympus). Line intensity profiles were produced using FLUOVIEW ver. 4.2 (Olympus).

Quantitative evaluation of temporal changes in localization of LZY4-mScarlet along the gravity direction

To quantitatively evaluate localization changes of the LZY4-mScarlet in the plasma membranes after the change in gravity direction, we measured the LZY4-mScarlet intensity weighted by the opposite direction of gravity. When LZY4-mScarlet is accumulated at a higher position, the weighted intensity shows a higher value. When LZY4-mScarlet is accumulated at a lower position, the weighted intensity indicates a lower value. The workflow for the image processing was shown in fig. S11. All image processing and measurement were performed using ImageJ software (https://imagej.nih.gov/ij/). To eliminate fluorescent signals derived from neighboring cells, we manually traced the plasma membranes as a line of interest (LOI), excluding adjacent regions in which brighter fluorescence was observed (fig. S11A). If the LOI is less than 66% of the relative height of the cell due to the influence of neighboring cells, it was excluded from the analysis. Cell regions were also manually segmented as a region of interest (ROI) to identify the cell height (fig. S11D). Next, the fluorescent intensity on LOI (fig. S11B) was averaged along the x-axis to obtain the average intensity distribution in the direction of gravity (fig. S11C). Based on the binary image with the ROI (fig. S11E), a cell height image was obtained with the x-axis maximum intensity projection (fig. S11F). The average intensity distribution image was masked with the cell height image (fig. S11G). To standardize the cell height, the image was transformed into 100 pixels in height. The above process was performed for all time frames, and the processed images were aligned along the x-axis direction according to time to make a kymograph showing the temporal change in the average intensity distribution along the gravity direction (fig. S11H). Then, the gradient image (fig. S12I), in which the intensity decreases linearly from 100 to 1 in the gravity direction, was multiplied to the kymograph (fig. S11J). Finally, the time evolution of the intensity weighted by the opposite direction of gravity was obtained by integrating the weighted kymograph in the y-axis direction (Fig. 3, and fig. S11K).

Analyses with primary roots

To perform live cell imaging of *LZY4p::LZY4-mScarlet / lzy4* in primary roots, vernalized seeds were germinated in 1/2 MS liquid medium within a 2-well chambered coverglass (Nunc Lab-Tek, Thermo Fisher Scientific), where a plastic coverslip (Celldesk LF2, Sumitomo Bakelite) and pieces of 0.3-mm-thick silicone-rubber sheet as spacer were stacked on the coverglass to make space for root growing. The chambers were leaned against a wall of a moist box, allowing the roots to grow along the surface of the coverslip, and they were kept for 4 days. Then, the micro-chamber was launched on the vertical stage confocal microscope (*35*). This made it possible to detect faint

*LZY4p::LZY4-mScarlet* fluorescence in primary roots. LOIs were determined as indicated in fig. S12C, the ratio (b/a) of fluorescent intensity was calculated. For measurements of R2D2 and gravitropism, 4-day-old seedling of *LZY4p::LZY4-mScarlet / lzy4* was transferred to a chamber slide (NuncLab-Tek II Chambered Coverglass, Thermo Fisher Scientific) with surrounding agar medium (*59*). After 3-hrs incubation vertically in the growth chamber, the chamber slide was launched on the vertical stage confocal microscope. For R2D2 measurements, images were taken with the vertical stage confocal microscope (*35*) with 20x objective lens (UPlanApo 20x, 0.7NA, Olympus). DII-n3xVenus and mDII-ntdTomato were excited with 488 nm and 561 nm lasers and detected via 525/50 nm and 617/73 nm emission filters (Semrock), respectively. After gravistimulation by 90° degree-rotation of the rotatable stage, images were taken in 2 min-interval for 1 hr. The first 12 to 18 epidermal cells were used for measurements. ROIs containing the same number of epidermal cells were set on the upper and lower sides based on the images of mDII-ntdTomato, and the average values of fluorescence intensity of DII-n3xVenus and mDII-ntdTomato in the ROIs were measured. The ratio (DII/mDII) was calculated and then the ratio (Upper/Lower) was indicated in fig. S12E. MetaMorph (Molecular Device) and ImageJ (https://imagej.nih.gov/ij/) were used for image acquisition and analyses. For measurements of gravitropic response of primary roots, images were taken with an inverted microscope (ECLIPSE Ti2-E, Nikon) tilted by 90° (*59*) equipped with 20x objective lens (CFI Plan Fluor 20x, 0.5 NA, Nikon) and Zyla sCMOS (ANDOR). NIS-Elements RA (Nikon) and ImageJ were used for image acquisition and analyses. By subtracting the root tip angles at Time 0 from those of each time point, curvature (degree) was indicated in fig. S12D.

Photoconversion of mEos2

To analyze the subcellular dynamics of LZY4 in LR columella cells, the amyloplasts on which LZY4-mEos are located were irradiated with 405-nm light (laser power, 20% of 12 mW; 4 repeats of 1s pulse with 1 s interval) of a patterned illumination device (Pixel illuminator; Pinpoint Photonics, inc, Yokohama) equipped on the vertical-stage confocal microscope, and we monitored the red fluorescence of mEos2 on the plasma membrane proximal to the amyloplasts after photoconversion.

For quantification of the fluorescent changes, the Line of Interest (LOI) with 3-pixel-width was drawn orthogonally to the plasma membrane region proximal to the amyloplasts, and the intensity profile on the LOI was obtained. The values at the positions intersecting with the plasma membrane were used as the fluorescent intensities on the plasma membrane. The positions were determined by referring to the bright field images. The background values were subtracted from the original fluorescence images by "Subtract Background" (rolling ball radius = 15 pixels) of ImageJ (https://imagej.nih.gov/ij/) before quantification.

LZY4-mScarlet dynamics during amyloplast manipulation

Surface-sterilized seeds were sown under a thin layer of MS medium solidified on a cover slip (24 x 50 mm) in a plate, and cultivated under a long-day cycle of 16h light/ 8 h darkness at 22°C in a growth chamber. The plates were leaned against a wall of the growth chamber, allowing the roots to grow between the medium and cover slip. Young lateral roots with a length of < 3 mm were used for the following experiments.

Amyloplasts were remotely manipulated with optical tweezers as previously described (*41*). Briefly, a near-infrared laser beam ($\lambda$ = 1064 nm) from a continuous wave $Nd{:}YVO_4$ laser (BL-106SU-FE, Spectra-Physics) was introduced into an inverted confocal laser scanning microscope

(A1R MP+, Nikon), and focused on the root cells via an oil immersion objective lens (NA = 1.35; Olympus). The power of a laser for optical tweezer was measured with a thermal sensor (12A-V1-ROHS, Ophir) equipped with a laser power meter display (VEGA, Ophir) and adjusted to be less than 10 mW with a half-wave plate and a polarizing prism.

The fluorescence and bright-field images were acquired with the confocal laser scanning microscope and imaging software (NIS-Elements Confocal, Nikon). mScarlet was excited with a 561 nm laser and the emitted fluorescence passing through a 595/50 nm filter was obtained. To monitor the localization of LZY4-mScarlet in real-time (Fig. 4C), we moved amyloplasts toward the plasma membrane using optical tweezers and analyzed changes in the LZY4-mScarlet fluorescence intensity at regions of interest (ROIs) set on the plasma membrane using an image analysis software (NIS-Elements Advanced Research Analysis, Nikon). The fractional fluorescence changes ($\Delta F/F_0$) were calculated over time using the equation $\Delta F/F_0 = (F - F_0)/F_0$ (*60*) where F and $F_0$ indicate the fluorescence intensity of LZY4-mScarlet at an arbitrary time and the averaged baseline fluorescence defined by the average of F over the first 3 frames of the recording, respectively. We also acquired z-stack images (5 µm in depth, 0.5 µm step) of LZY4-mScarlet before and after moving amyloplasts in ~3 min (figs. S15 and S16). The z-stack images were converted into maximum intensity projection images. ROIs were set on the lower and upper sides of the plasma membrane of the columella cells. Each fluorescence intensity was divided by the averaged fluorescence intensity between the lower and upper side of the plasma membrane and the ratio of the fluorescence intensity before and after the amyloplast manipulation was calculated. For control experiments, the trapping laser was irradiated at the plasma membrane for ~1 min without manipulating amyloplasts. To avoid photobleaching of LZY4-mScarlet during manipulating amyloplasts, the positions of the trapped amyloplasts and the plasma membrane were confirmed using bright-field images visualized with a 639 nm laser. Changes in LZY4-mScarlet localization in response to the amyloplast movements were analyzed by unpaired *t*-test with Welch's correction using GraphPad Prism 7 (GraphPad Software).

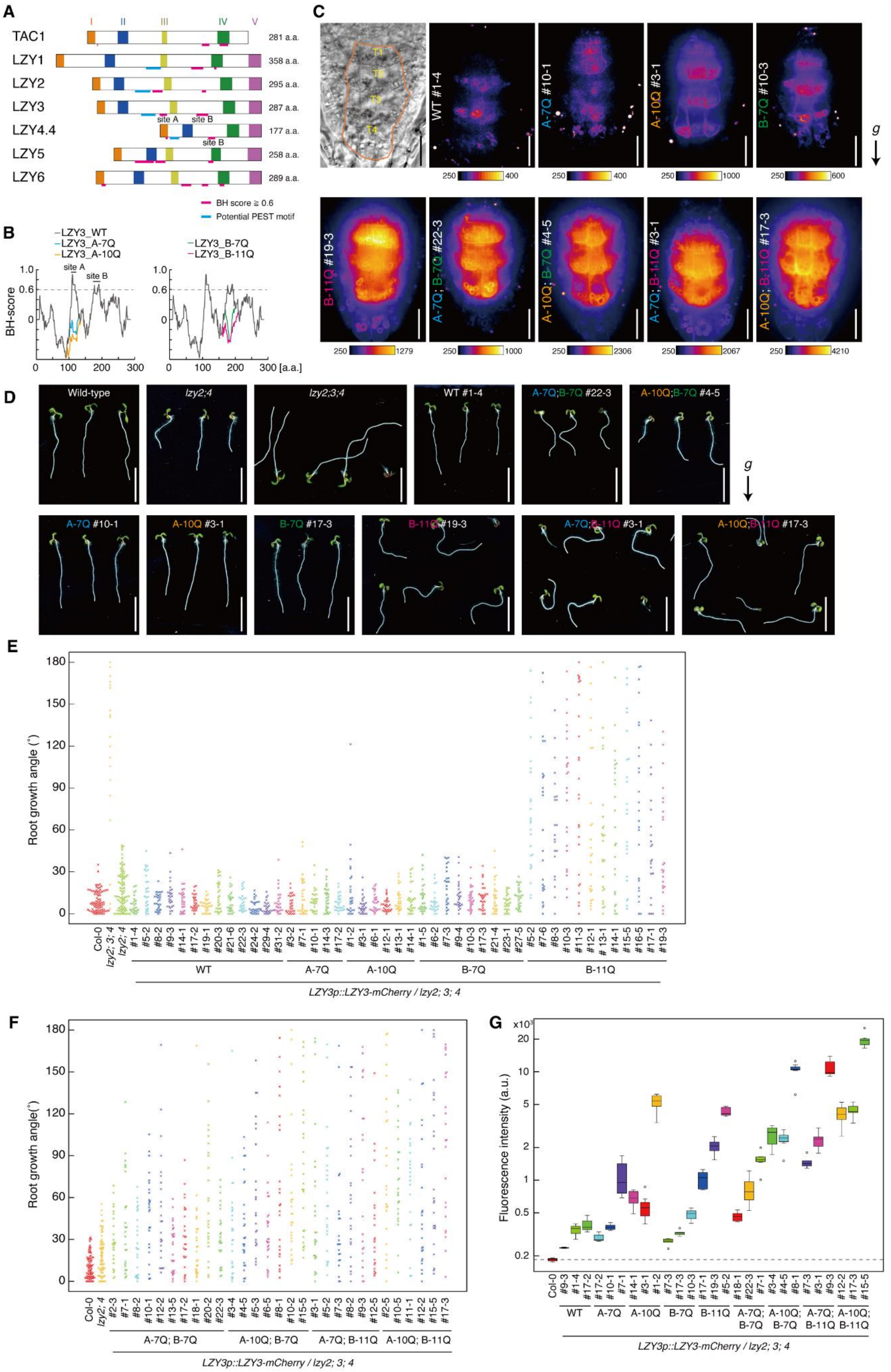
A
TAC1
LZY1
LZY2
LZY3
LZY4.4
LZY5
LZY6
I
II
III
IV
V
281 a.a.
358 a.a.
295 a.a.
287 a.a.
177 a.a.
258 a.a.
289 a.a.
site A
site B
BH score ≧ 0.6
Potential PEST motif
B
LZY3_WT
LZY3_A-7Q
LZY3_A-10Q
LZY3_B-7Q
LZY3_B-11Q
BH-score
[a.a.]
C
T1
T2
T3
T4
WT #1-4
A-7Q #10-1
A-10Q #3-1
B-7Q #10-3
B-11Q #19-3
A-7Q; B-7Q #22-3
A-10Q; B-7Q #4-5
A-7Q; B-11Q #3-1
A-10Q; B-11Q #17-3
g
D
Wild-type
lzy2;4
lzy2;3;4
WT #1-4
A-7Q;B-7Q #22-3
A-10Q;B-7Q #4-5
A-7Q #10-1
A-10Q #3-1
B-7Q #17-3
B-11Q #19-3
A-7Q;B-11Q #3-1
A-10Q;B-11Q #17-3
E
Root growth angle (°)
Col-0
lzy2; 3; 4
lzy2; 4
WT
A-7Q
A-10Q
B-7Q
B-11Q
LZY3p::LZY3-mCherry / lzy2; 3; 4
F
Root growth angle(°)
A-7Q; B-7Q
A-10Q; B-7Q
A-7Q; B-11Q
A-10Q; B-11Q
LZY3p::LZY3-mCherry / lzy2; 3; 4
G
Fluorescence intensity (a.u.)
WT
A-7Q
A-10Q
B-7Q
B-11Q
A-7Q; B-7Q
A-10Q; B-7Q
A-7Q; B-11Q
A-10Q; B-11Q
LZY3p::LZY3-mCherry / lzy2; 3; 4

**Fig. S1.**

Potential membrane association sites in LZY proteins and their function. (**A**) Schematic structure of Arabidopsis LAZY1-LIKE family proteins. Five conserved regions (I-V) are indicated as colored boxes (*10*). The sites that show BH-scores ≥ 0.6, and potential PEST motifs are displayed as magenta and cyan bars, respectively, attached to each structure. The positions of potential membrane association sites A and B for LZY3 or site B for LZY4 are labeled near the bars. (**B**) BH-score distribution throughout the full length of LZY3 protein shows two peaks above the threshold value 0.6 (dashed line), which predicts membrane association sites. The amino acid substitutions at sites A and B affect the BH-score around each site, respectively. (**C**) Representative confocal images of wild-type (WT), and all the generated mutants on site A and B of LZY3-mCherry in columella cells of primary roots. The seedlings were kept vertically before and during the imaging. The roots of B-11Q, A7Q;B-7Q, A-10Q;B-7Q, A-7Q;B-11Q, and A-10Q;B-11Q lines were straightened at least 3 hours before imaging to make the amyloplasts sedimented toward the root tip, because those roots were meandering as shown in **D**. All images were acquired with an identical optical setting, and the color-coded heatmaps for signal intensities are placed below each image. Scale bars, 20 µm. *g*, the direction of gravity. (**D**) Representative images of 5-day-old seedlings of wild-type, *lzy2;4*, *lzy2;3;4*, and transgenic plants expressing LZY3-mCherry mutants on the site A and/or B, which are used in **C**, **E**, **F, G** and Fig. 1. The expression of the transgenes were driven by *LZY3* promoter. Scale bars, 1 cm. (**E**) Growth angle of primary root tips of 5-day-old seedlings of wild-type, *lzy2;4*, *lzy2;3;4*, and transgenic plants expressing LZY3-mCherry mutants on the site A or B. (**F**) Growth angle of primary root tips of 5-day-old seedlings of wild-type, *lzy2;4*, and transgenic plants expressing LZY3-mCherry multiple mutants on the site A and B. Direction of each root tip was measured as the absolute value toward the direction of gravity. **(G)** Protein accumulation levels of LZY3-mCherry in columella cells of primary roots. Mean values of the fluorescence intensity in the area including tier 1 to 3 cells were quantified for the representative lines (3 to 8 roots) and plotted as boxplots. The y-axis is represented as a logarithmic scale. Center lines show the medians; box limits indicate the 25th and 75th; whiskers extend 1.5 times the interquartile range from the 25th and 75th percentiles; outliers are represented by dots. The dashed line extended from the Col-0 value means the background signal.

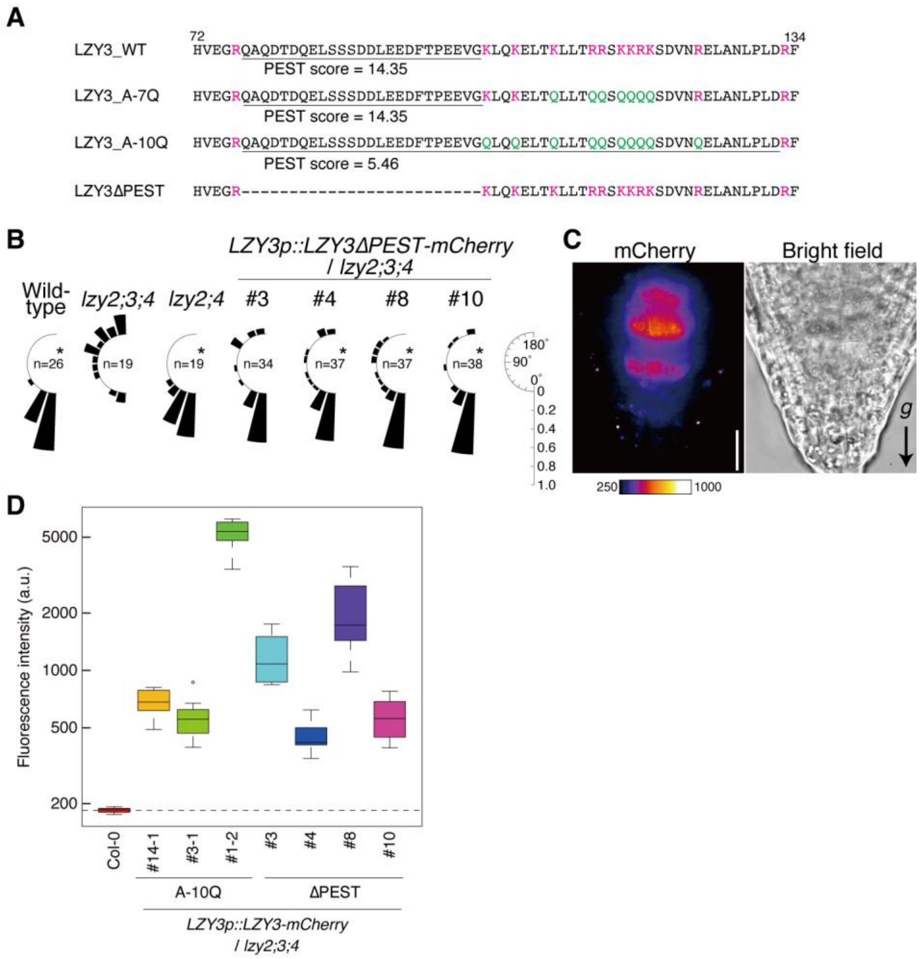


**Fig. S2.**

(**A**) Sequences containing a potential PEST motif and site A in LZY3 and its mutant proteins. The PEST motif was predicted by the algorithm epestfind (https://emboss.bioinformatics.nl/cgi-bin/emboss/epestfind) and the calculated scores were shown. The score of LZY3_A-10Q is largely reduced by substitutions of basic residues by glutamine residues, although the predicted region is expanded. Deletion of the PEST motif does not show a significant score (LZY3ΔPEST). (**B**) Root growth angle of wild-type, *lzy2;3;4*, *lzy2;4*, and transgenic plants expressing LZY3 mutant protein which lacks the PEST motif under the control of *LZY3* promoter in *lzy2;3;4* mutant background (T2 generations). The direction of each root tip was measured as the absolute value toward the direction of gravity. The frequency was calculated as the proportion of root numbers that fall within intervals of 15˚ to the total number of the analyzed roots for each line (range, 0 to 180˚). The chart of *lzy2;3;4* is identical to Fig. 1. (**C**) A representative confocal image of LZY3ΔPEST-mCherry (line #4) in columella cells of primary roots. The seedlings were kept vertically before and during the imaging. The color-coded heatmap for signal intensities is placed below the image. Scale bars, 20 µm. *g*, the direction of gravity. (**D**) Protein accumulation levels of LZY3-mCherry in columella cells of primary roots. Mean values of the fluorescence intensity in the area including tier 1 to 3 cells were quantified for the representative lines (6-8 roots) and plotted as boxplots like fig. S1G. The plots on Col-0 and A-10Q are identical to fig. S1G.

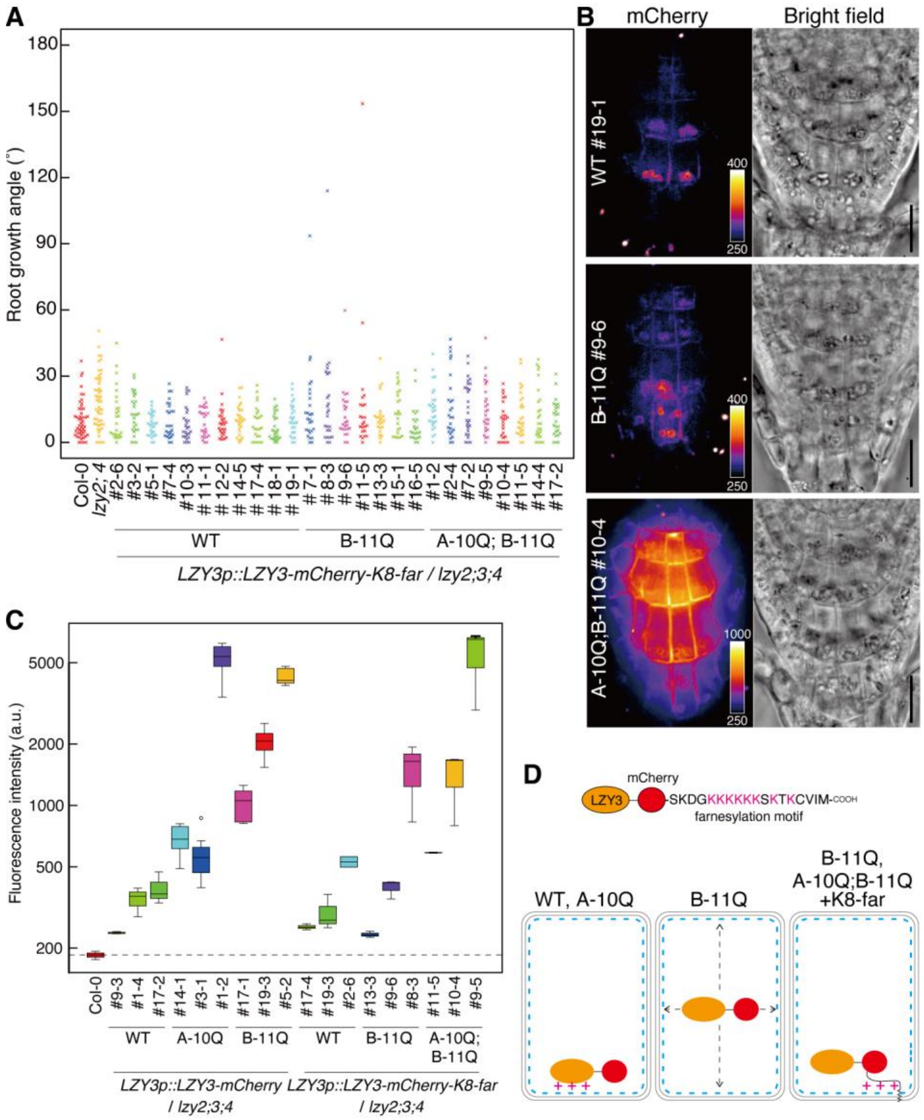


**Fig. S3.**

Addition of a farnesylation motif recovers the impaired root gravitropic phenotype of transgenic plants with LZY3-mCherry harboring B-11Q and A-10Q;B-11Q mutations. (**A**) Growth angle of primary root tips of 5-day-old seedlings of wild-type, *lzy2;4*, and transgenic plants expressing LZY3-mCherry with a farnesylation motif from K-Ras (*48*) at the C-terminus of mCherry (LZY3-mCherry-K8-far) (see **D**) into *lzy2;3;4* background. The expression of LZY3-mCherry-K8-far proteins that harbor B-11Q, A-10Q;B-11Q, or no mutations were driven by *LZY3* promoter. Direction of each root tip was measured as the absolute value toward the direction of gravity. (**B**) Representative confocal images of LZY3-mCherry-K8-far harboring no substitution, B-11Q, and A-10Q;B-11Q in columella cells of primary roots. The seedlings were kept vertically before and during the imaging. The color-coded heatmaps for signal intensities are placed within each image. Scale bars, 20 µm. **(C)** Protein accumulation levels of LZY3-mCherry in columella cells of primary roots. Mean values of the fluorescence intensity in the area including tier 1 to 3 cells were quantified for the representative lines (2-8 roots) and shown as boxplots like fig. S1G. The plots on Col-0, and LZY3_WT/A-10Q/B-11Q are identical to fig. S1G. (**D**) Illustration of LZY3-

mCherry-K8-far and a summary of LZY3 recruitment to the plasma membrane. WT and A-10Q of LZY3-mCherry have positive charges on their protein surfaces, which could be sufficient for electrostatic interaction with anionic lipids distributed on the plasma membrane (also see fig. S13). In the B-11Q protein, the reduced positive charge could not be enough for the tight association with the plasma membrane to transduce the gravity signals precisely, although the weak association was still observed (Fig. 1, and fig. S1). Addition of the farnesylation motif recovers the recruitment of LZY3-mCherry harboring B-11Q and A-10Q:B-11Q to the plasma membrane via the electrostatic interaction and acylation anchor.

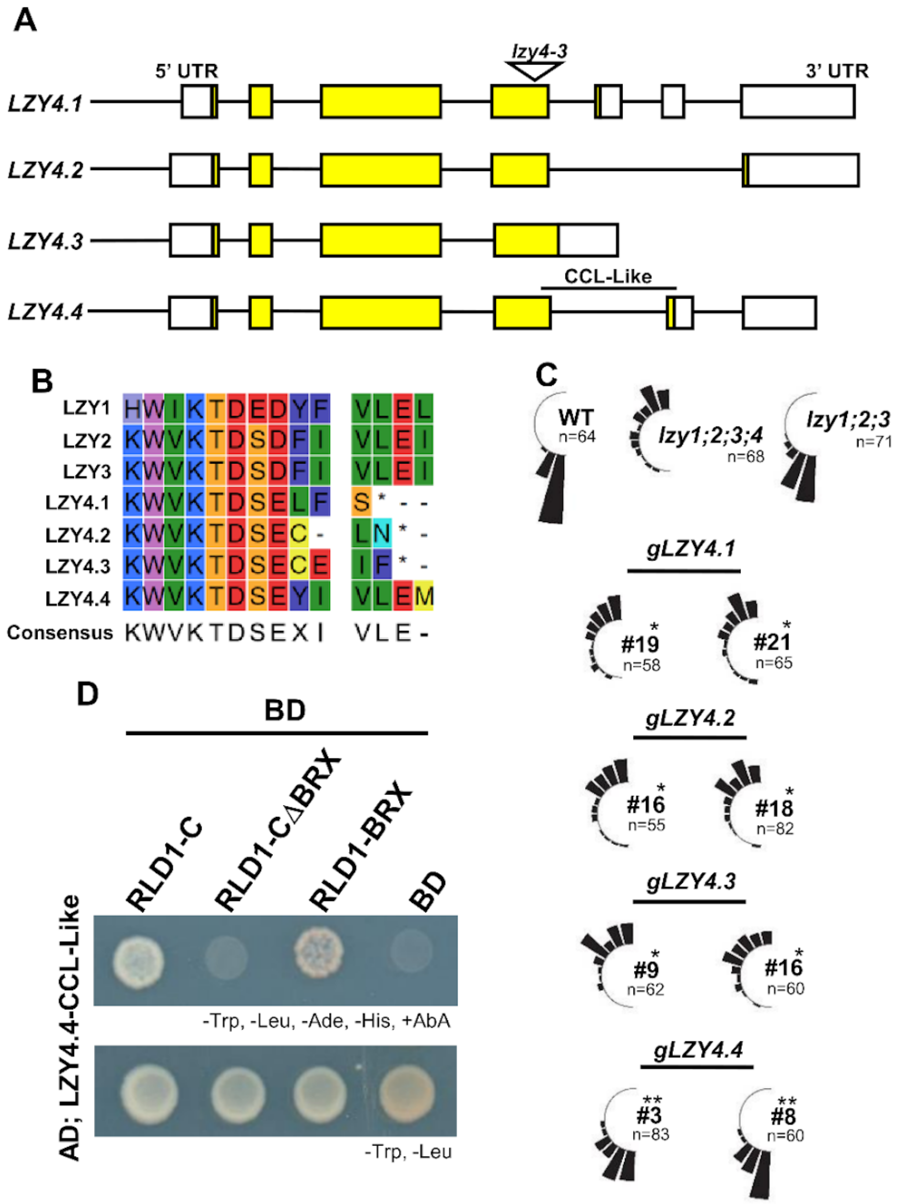


**Fig. S4.**

LZY4.4 functions in the root gravitropic response. **(A)** Splicing variants of *LZY4*. White squares indicate UTRs; yellow squares indicate CDS (Gene bank accession *LZY4.1*; NM_001123839, *LZY4.2*; NM_001123840, *LZY4.3*; NM_001123841, *LZY4.4*; NM_001332385). *lzy4-3* (GABI_479C08) was used in this study as *lzy4* mutant. **(B)** Comparison of C-terminal sequences among *LZY* genes. The CCL-Like sequence is found only in LZY4.4 among the splicing variants. **(C)** Complementation test for *LZY4* splicing variants. Constructs expressing the splicing variant of *LZY4*, respectively, were transformed into *lzy1;2;3;4*, and orientation of the primary roots was measured. Only the construct expressing *LZY4.4*, in which the CCL-Like sequence is conserved, complemented the *lzy1;2;3;4* phenotype. Asterisks indicate significant differences [$P < 0.0045$ (0.05/11)] by Wilcoxon's rank sum test with Bonferroni's correction (*, compared with *lzy1;2;3*; **, compared with *lzy1;2;3;4*). **(D)** Interaction between CCL-Like of LZY4.4 and C-terminally truncated forms of RLD1 in the Y2H system.

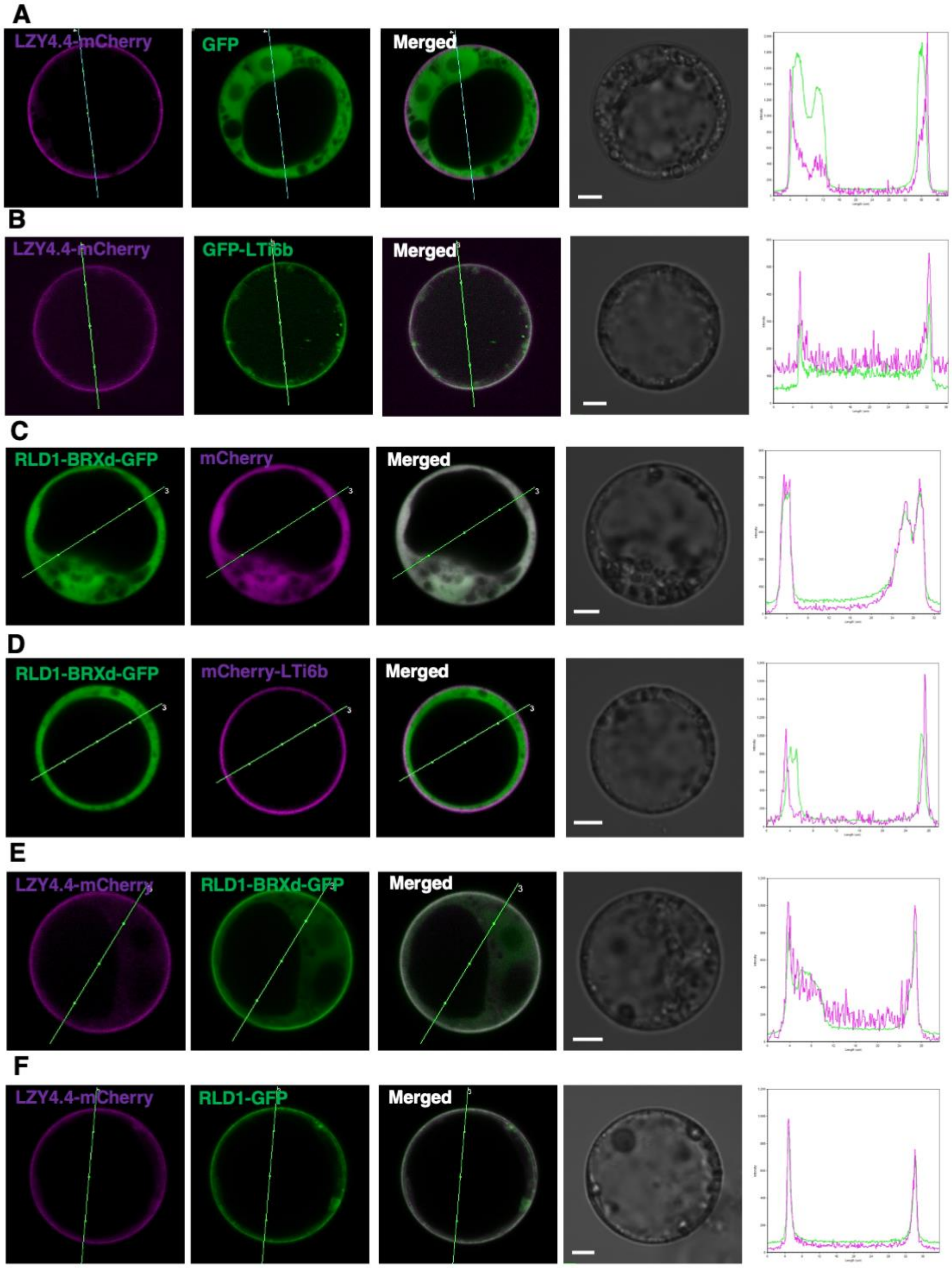


**Fig. S5.**

Transient expression assay in protoplasts of cultured Arabidopsis cells. GFP and mCherry signals in regions of interest (lines) were shown in line intensity profiles. Scale bars, 5 µm. **(A)** Co-expression of LZY4.4-mCherry with GFP. **(B)** Co-expression of LZY4.4-mCherry with GFP-LTI6b. **(C)** Co-expression of RLD1-BRXdomain-GFP (RLD1-BRXd-GFP) with mCherry. **(D)** Co-expression of RLD1-BRXd-GFP with mCherry-LTI6b. **(E)** Co-expression of LZY4.4-mCherry with RLD1-BRXd-GFP. **(F)** Co-expression of LZY4.4-mCherry with RLD1-GFP.

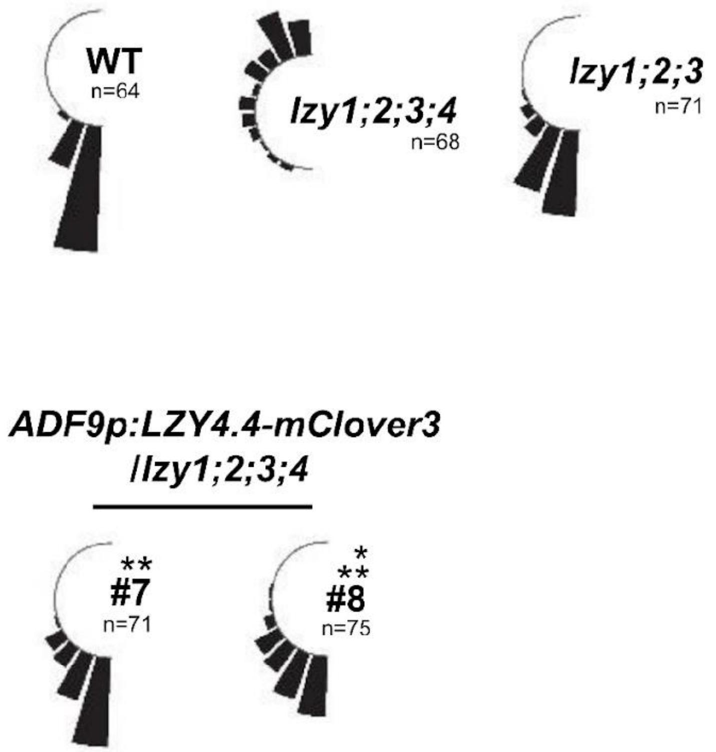


**Fig. S6.**

Complementation test for *LZY4.4* using the statocyte-specific *ADF9* promoter. The CDS of *LZY4.4-mClover3* was expressed in *lzy1;2;3;4* background under the control of the *ADF9* promoter and the direction of the primary roots was measured. Asterisks indicate significant differences [$P$ < 0.01 (0.05/5)] by Wilcoxon's rank sum test with Bonferroni's correction (*, compared with *lzy1;2;3*; **, compared with *lzy1;2;3;4*).

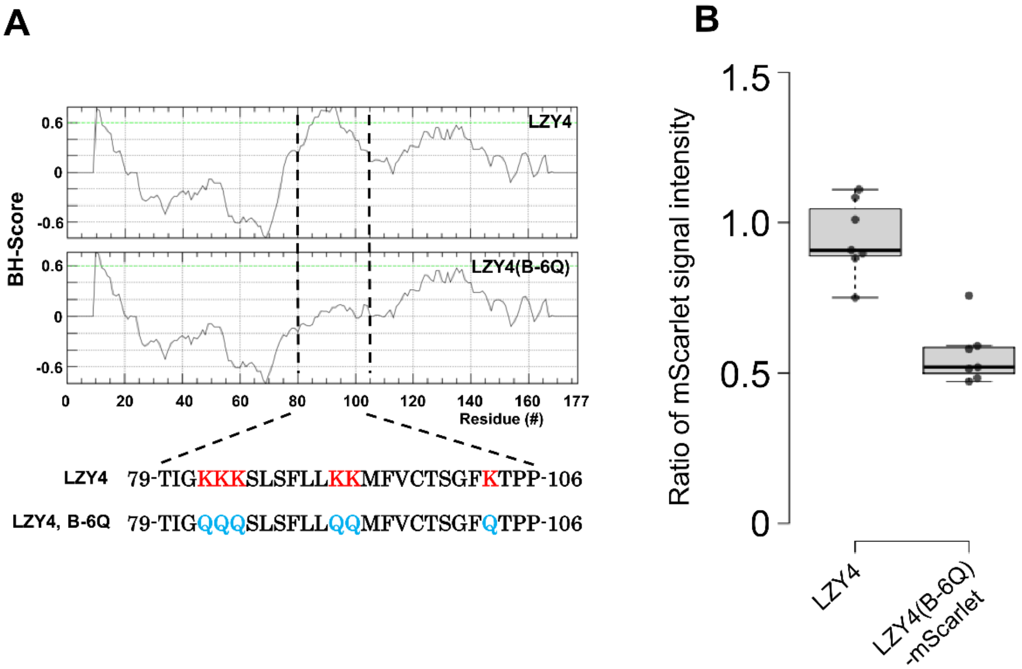


**Fig. S7.**

Characterization of LZY4(B-6Q)-mScarlet. **(A)** BH score distribution throughout the full length of LZY4 and LZY4(B-6Q). Lysine 83-85, 92 and 93, and 102 of LZY4 were replaced with glutamine. **(B)** Comparison of fluorescence signal intensities between LZY4-mScarlet and LZY4(B-6Q)-mScarlet on the plasma membrane. Signal values on the amyloplast and on the plasma membrane were determined using ImageJ. The ratio of the signal intensities was calculated (plasma membrane/amyloplasts) and shown in a boxplot (n = 7). Center lines show the medians; box limits indicate the 25th and 75th; whiskers extend 1.5 times the interquartile range from the 25th and 75th percentiles; outliers are represented by dots; data points are plotted as open circles. Differences between the means were assessed for statistical significance using a Student *t*-test ($P < 0.001$).

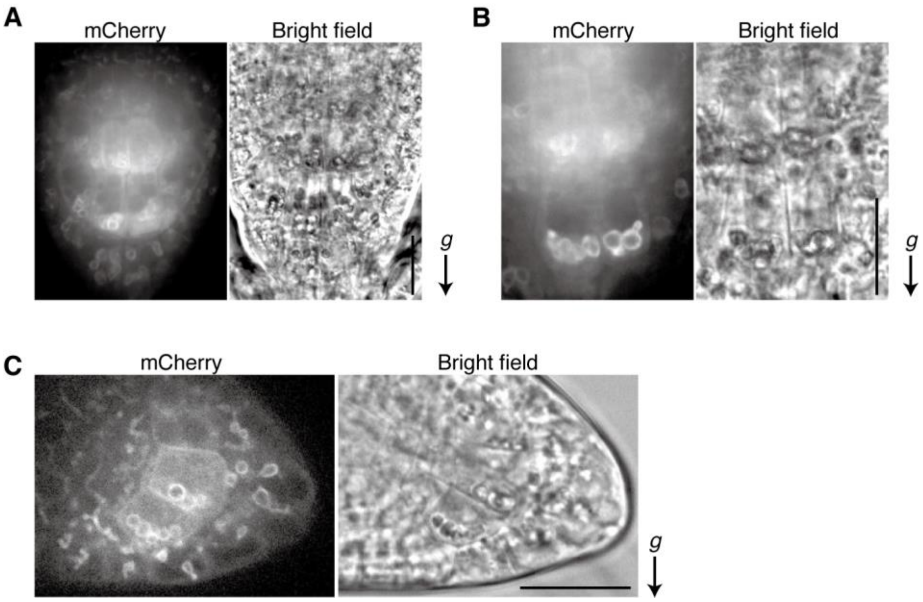


**Fig. S8.**

An N-terminal region of LZY3 is sufficient for localization on the amyloplast. An N-terminal region, 1-54 amino acid, of LZY3 was fused to the N-terminus of mCherry and expressed under the control of *ADF9* promoter in wild-type background (*ADF9p::LZY3(1-54)-mCherry*). The primary root tips of 5-day-old seedlings **(A** and **B)**, and the lateral root tips of 7-day-old seedlings **(C)** were observed (line #3, T2 generations). The strong fluorescent signals on the amyloplasts and faint signals on the plasma membrane were detected. In **C**, the plasma membrane signals did not show polarity toward gravity. Scale bars, 20 µm. *g*, the direction of gravity.

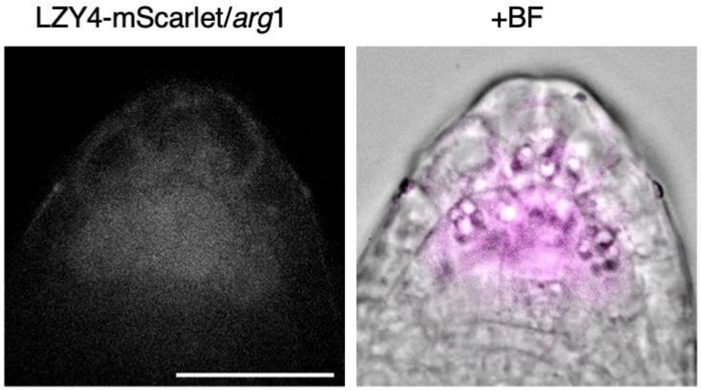


**Fig. S9.**

Subcellular localization of LZY4-mScarlet in columella cells of a lateral root of the *arg1* mutant. Scale bars, 25 µm.

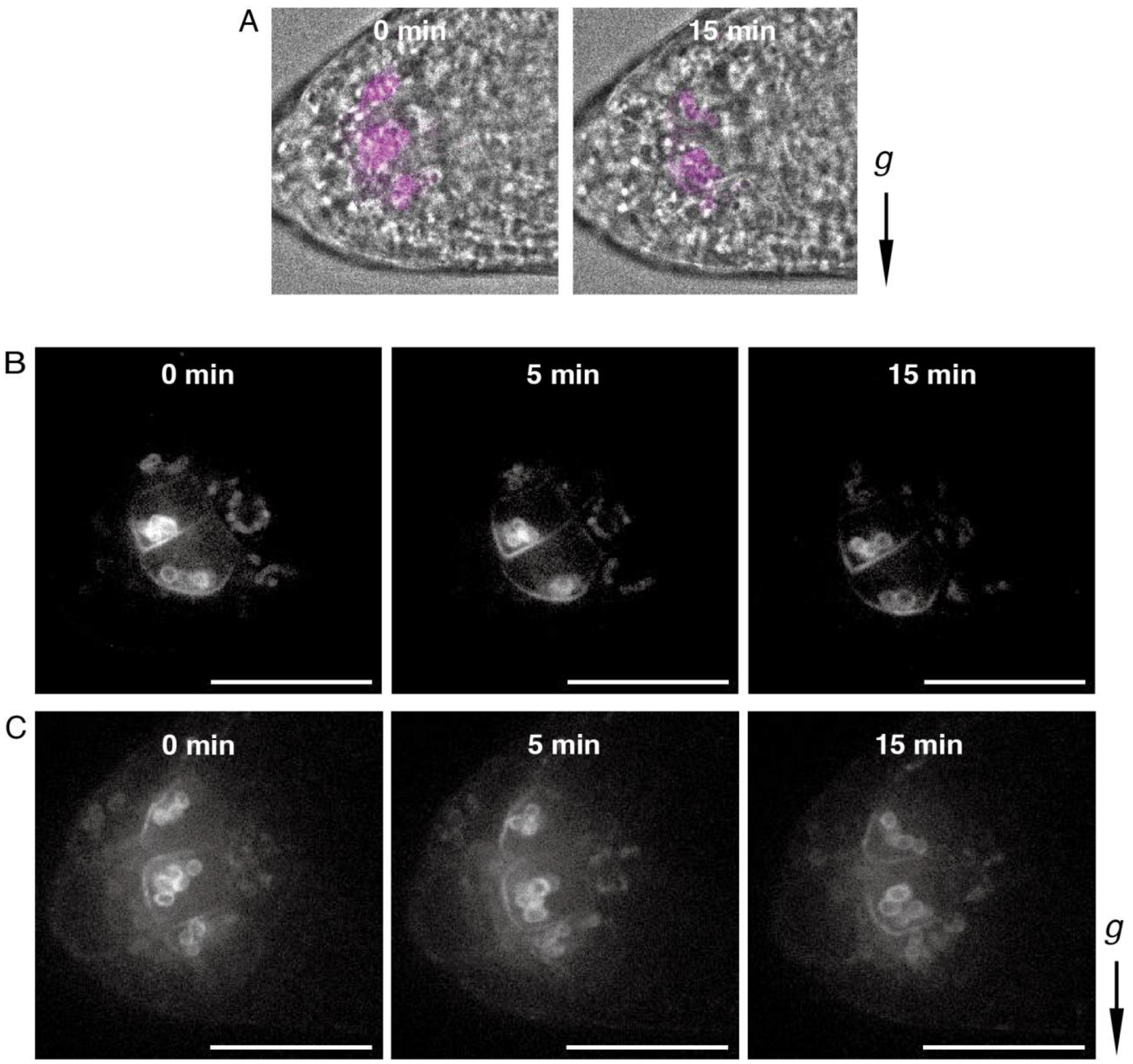


**Fig. S10.**

Polar localization of LZY4-mScarlet on the PM. **(A)** Representative merged images of fluorescence and bright field, corresponding to Fig. 3B. Stage 2 (< 3 mm) LR tips of *LZY4p::LZY4-mScarlet* / *lzy4* were observed. **(B** and **C)** Representative images for temporal changes of LZY4-mScarlet without **(B)** and with **(C)** gravistimulation. *g*, the direction of gravity. Scale bars, 25 µm.

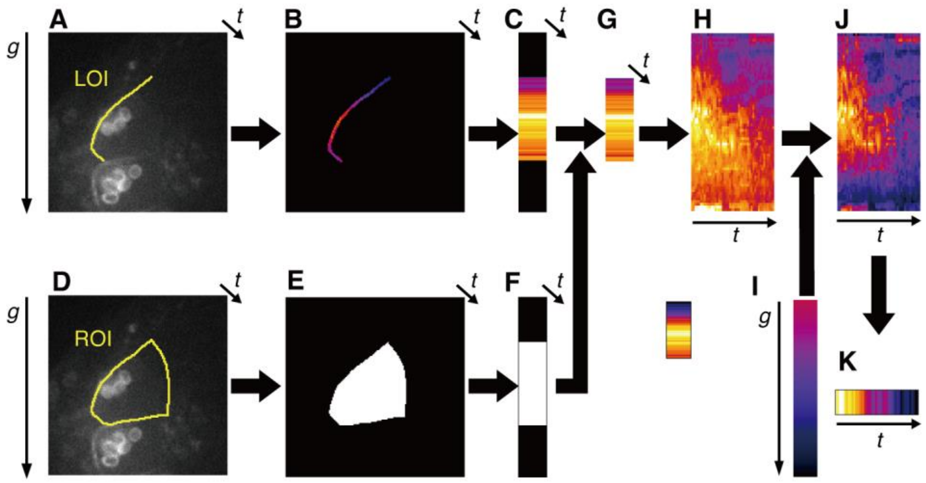


**Fig. S11.**

Workflow of the image processing to measure the LZY4-mScarlet intensity weighted by the opposite direction of gravity. **(A)** Line of interest (LOI) manually set on the plasma membranes in the time-lapse images. Arrow indicates the direction of gravity. **(B)** LZY4-mScarlet intensity in the LOI in image **(A)** visualized by pseudo color. **(C)** Average intensity image along the x-axis of **(B)**. **(D)** Region of interest (ROI) of the cell that was manually set. **(E)** Mask image of the cell generated with ROI in **(D)**. **(F)** Cell height image that is maximum intensity projection along the x-axis in **(E)**. **(G)** Masked average intensity image along the x-axis. The cell height was standardized into 100 pixels. **(H)** Kymograph of the average intensity distribution in the direction of gravity on LOI. **(I)** Gradient image for weighting by the position in opposite direction of the gravity. **(J)** Distribution of the intensity weighted by the opposite direction of gravity. **(K)** Time evolution of the intensity weighted by the opposite direction of gravity.

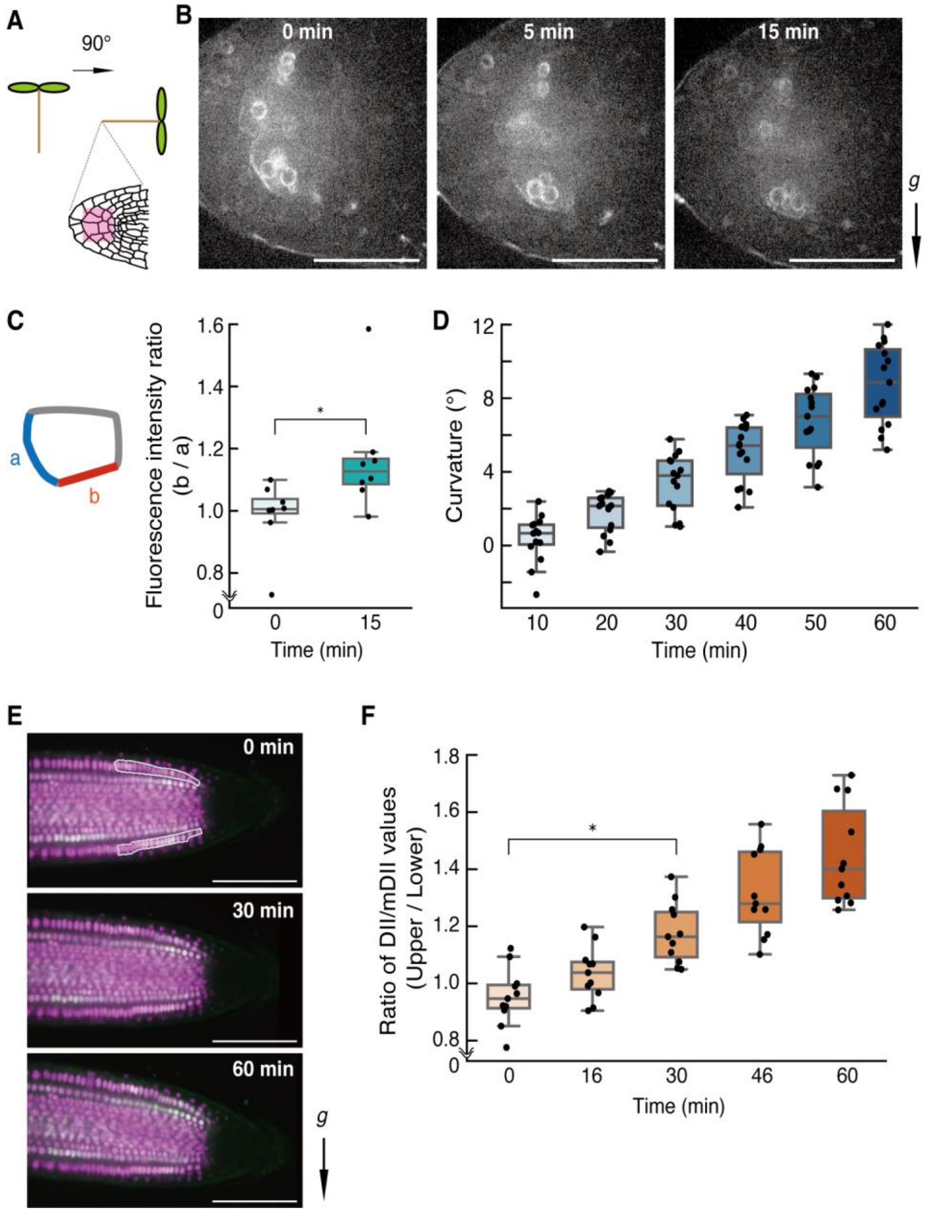


**Fig. S12.**

Temporal relationship among LZY4-mScarlet repolarization **(B and C)**, gravitropic curvature **(D)**, and formation of asymmetric auxin distribution **(E and F)** in the primary root. **(A)** Gravistimulation and site of observation. **(B)** A four-day-old seedling of *LZY4p::LZY4-mScarlet* / *lzy4* grown in a hydroponic micro-chamber was gravistimulated by 90° inclination. mScarlet was excited at 561 nm and the fluorescence passed through a 617/73 nm emission filter was detected. The images were taken at 0 min, 5 min, and 15 min after gravistimulation. Scale bars, 25 μm. **(C)** LZY4 repolarization was evaluated as the ratio of fluorescence intensities (b/a). LOIs were set on the PM lower side of the cell before (a) and after (b) gravistimulation. n = 8. Error bars, standard deviation (SD). * $P < 0.05$, Dunnett's test. **(D)** Gravitropic curvature of the primary roots grown in the chamber slide was measured. n = 8. Error bars, SD. **(E)** Auxin distribution in root tips was visualized by ratiometric imaging using R2D2. ROIs were set on the epidermis of the upper and lower sides. *g*, the direction of gravity. Scale bars, 100 μm. **(F)** DII/mDII values were calculated for the upper and lower sides, and the ratio of the values at the upper to that at the lower sides (Upper/Lower) was determined to evaluate auxin distribution in the root tip (n = 11). Error bars, SD. * $P < 0.05$, Dunnett's test.

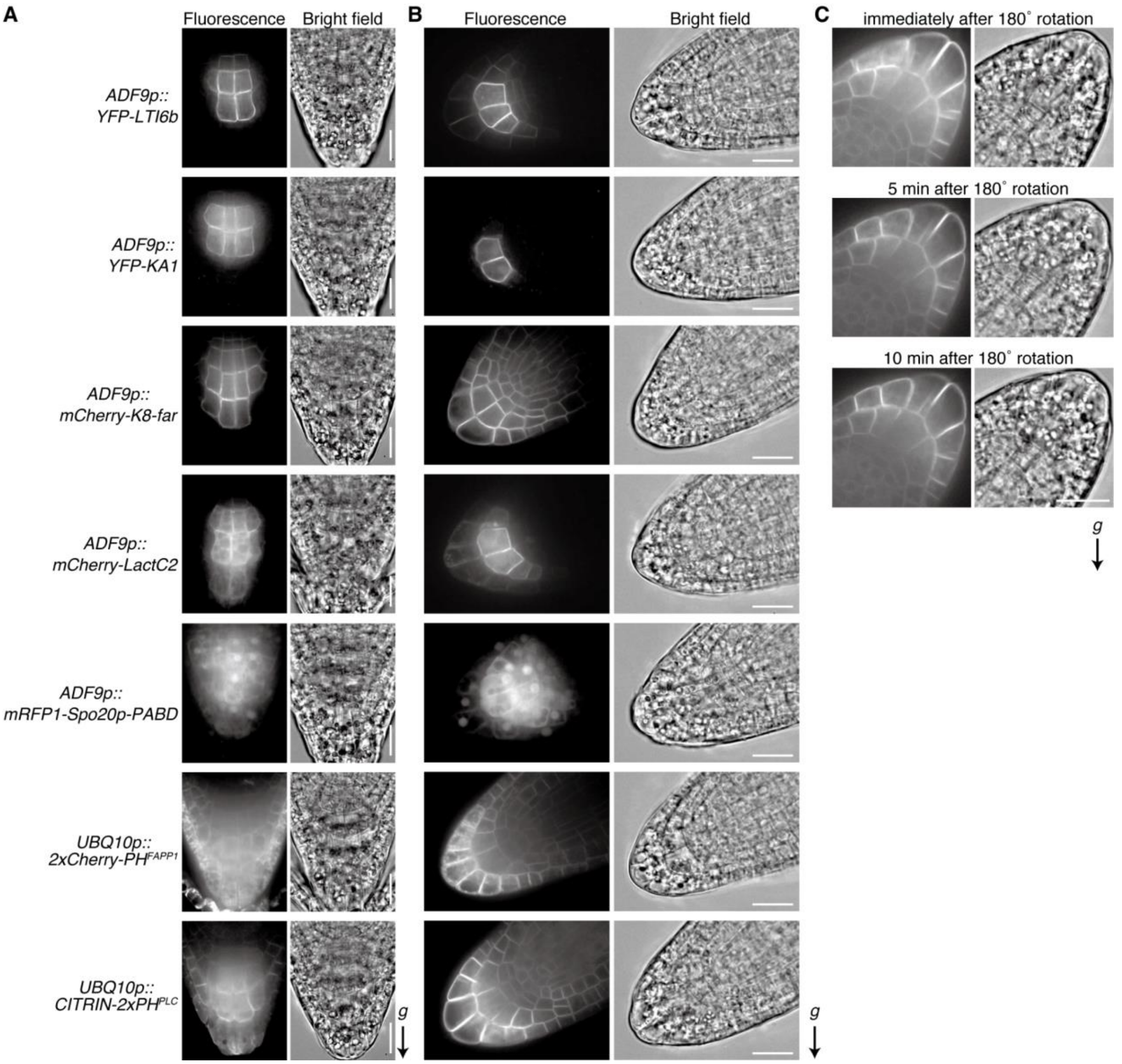


**Fig. S13.**

Distribution of anionic lipids in columella cells. Representative confocal images of the plasma membrane marker YFP-LTI6b, membrane surface charge reporters YFP-KA1$^{MARK1}$ and mCherry-K8-far, phosphatidylserine (PS)-biosensor mCherry-LactC2, phosphatidic acid (PA)-biosensor mRFP1-Spo20p-PABD, phosphatidylinositol-4-phosphate (PI(4)P)-biosensor 2xCherry-PH$^{FAPP1}$ (P5R), and phosphatidylinositol-4,5-bisphosphate (PI(4,5)$P_2$)-biosensor CITRIN-2xPH$^{PLC}$ (P24Y) in columella cells of primary roots of 5-day-old seedlings (**A**) and lateral roots of 7-day-old seedlings (**B**). Only CITRIN-2xPH$^{PLC}$ tends to be polarly distributed at the distal plasma membrane in columella cells of both primary and lateral roots. (**C**) A primary root expressing CITRIN-2xPH$^{PLC}$ was rotated 180˚ and time-lapse images were acquired on the same section for 10 min. The seedlings were kept vertically before and during the imaging. Scale bars, 20 µm. *g*, the direction of gravity.

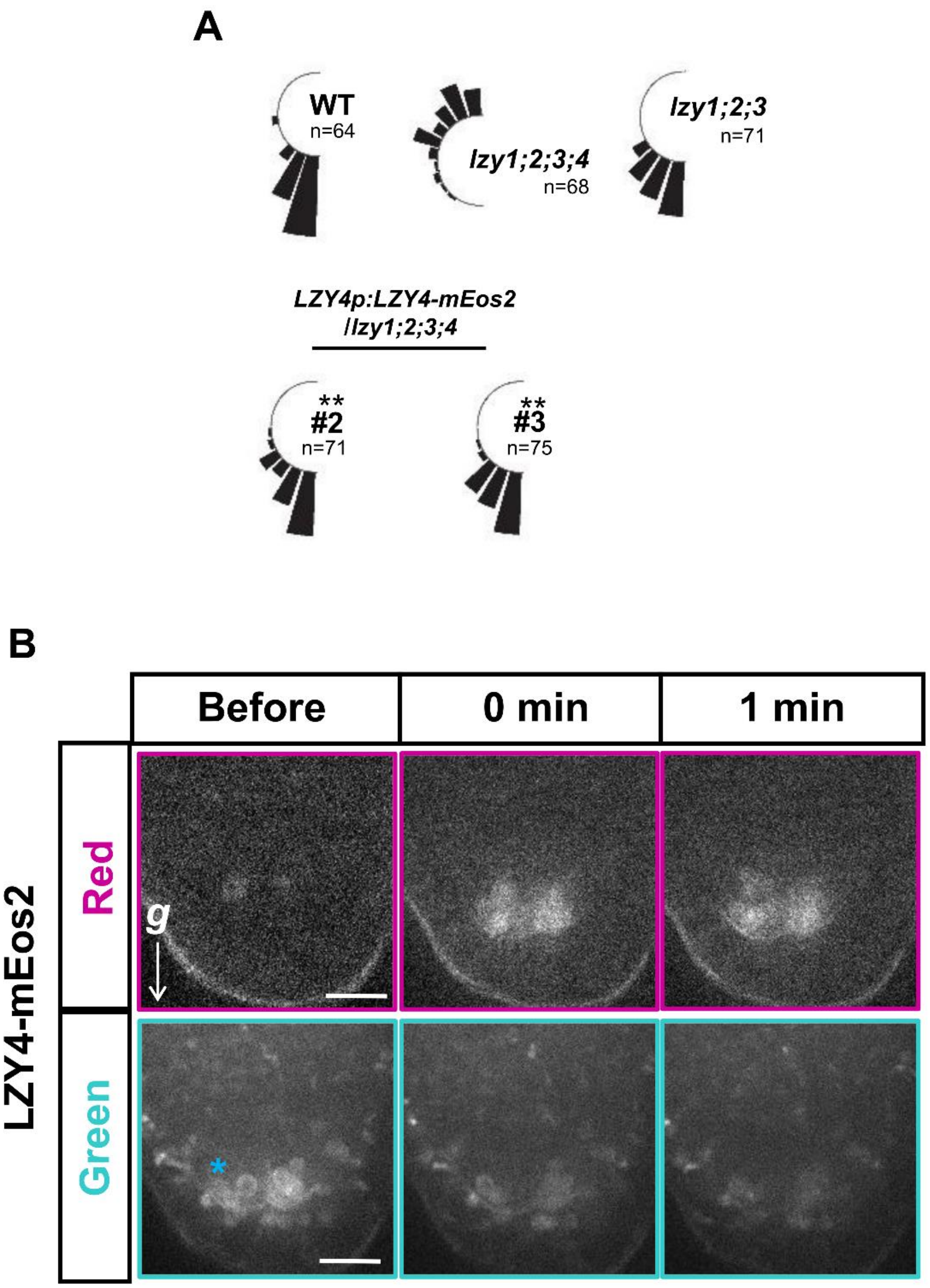


**Fig. S14.**

Photoconversion of LZY4-mEos2. **(A)** Complementation test of *LZY4p::LZY4-mEos2* / *lzy1;2;3;4* in the direction of primary root tip. Asterisks indicate significant differences [$P < 0.01$ (0.05/5)] by Wilcoxon's rank sum test with Bonferroni's correction (*, compared with *lzy1;2;3*; **, compared with *lzy1;2;3;4*). **(B)** LZY4-mEos2 (green) found in amyloplasts is photoconverted by laser irradiation. Point of interest, which is indicated as an asterisk in the bottom left image, was illuminated with a 405 nm laser. Scale bars, 10 µm. *g*, the direction of gravity.

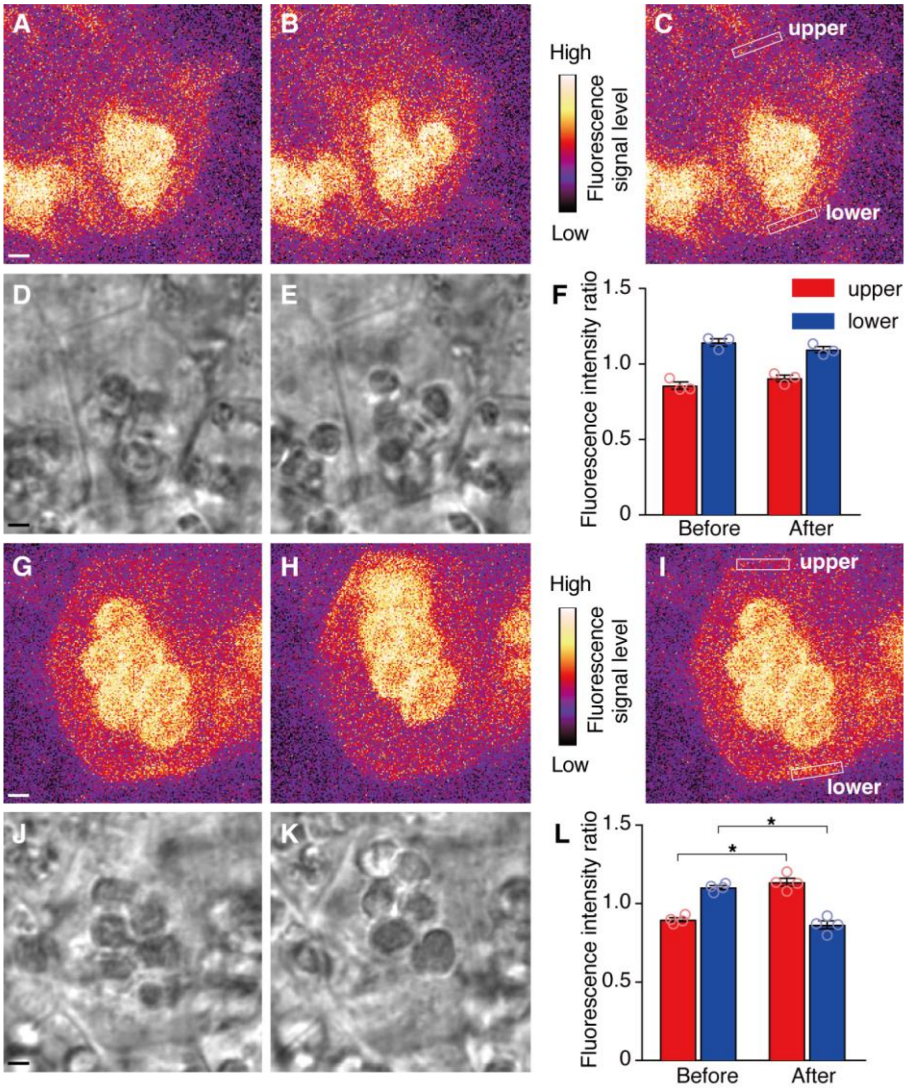


**Fig. S15.**

LZY4-mScarlet is repolarized according to amyloplast manipulation by optical tweezer. **(A** to **F)** Control experiment to examine the effect of irradiation with a laser for optical tweezer without manipulating amyloplasts on the LZY4-mScarlet behavior. Maximum Intensity Projection (MaxIP) image converted from Z-stack images of LZY4-mScarlet **(A** and **B)** and a representative optical slice of the bright-field images **(D** and **E)** before **(A** and **D)** and after **(B** and **E)** irradiation. **(C)** ROIs were set on the lower and upper sides of the PM of columella cells. **(F)** Each fluorescence intensity was divided by the averaged fluorescence intensity between the lower and upper sides of the PM, and the ratio of the fluorescence intensity before and after irradiation was calculated. Irradiation with a laser for optical tweezer had no effect on LZY4-mScarlet behavior. n = 3. **(G** to **L)** LZY4-mScarlet behavior upon amyloplast manipulation by optical tweezer. MaxIP image of LZY4-mScarlet **(G** and **H)** and bright-field images **(J** and **K)** before **(G** and **J)** and after **(H** and **K)** amyloplast manipulation. Trapped amyloplasts were moved to the upper side. **(I)** ROIs were set on the lower and upper sides of the PM of columella cells. **(L)** Ratio of the fluorescence intensity of the ROIs before and after irradiation was calculated as in **(F)**. n = 4. Scale bars, 2 μm. * $P < 0.001$, unpaired *t*-test with Welch's correction.

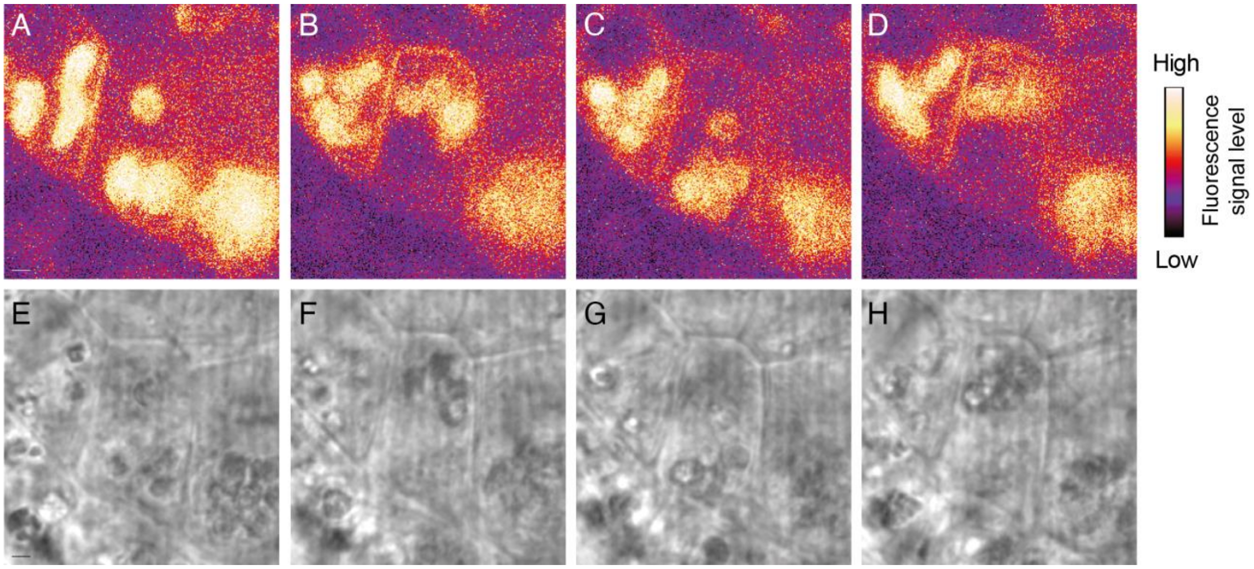


**Fig. S16.**

LZY4-mScarlet is repeatedly repolarized according to amyloplast manipulation by the optical tweezer. Amyloplasts were moved back and forth between the upper and lower of the cell. Maximum Intensity Projection (MaxIP) image converted from Z-stack images of LZY4-mScarlet (**A** to **D**) and bright-field images (**E** to **H**). (**A** and **E**) Most amyloplasts are positioned on the lower side of the cell before manipulation. The fluorescence signal of LZY4-mScarlet was observed at the PM of the lower side. (**B** and **F**) Amyloplasts were subsequently trapped and moved to the upper side of the cell. Fluorescence signals emerged at the upper side of the cell. (**C** and **G**) Amyloplasts were subsequently moved to the lower side of the cell. The fluorescent intensity of the upper side significantly decreased and that of the lower side seemed to increase. (**D** and **H**) Amyloplasts were trapped and moved to the upper side of the cell again. The fluorescent intensity of the upper side increased again. Each move took 3 min to operate.

**Movie S1.**

Columella cells of *LZY4p::LZY4-mScarlet / lzy4* young lateral root without gravistimulation. Time-lapse microscopy was performed on the vertical stage confocal microscope. Images were acquired at 20-second intervals for 15 min. *g*, the direction of gravity.

**Movie S2.**

Columella cells of *LZY4p::LZY4-mScarlet / lzy4* young lateral root with gravistimulation. Time-lapse microscopy was performed on the vertical stage confocal microscope. Images were acquired at 20-second intervals for 15 min. *g*, the direction of gravity. Image acquisition was initiated immediately after gravistimulation by rotating the stage 135 degrees.

**Movie S3.**

Columella cells of *LZY4p::LZY4-mScarlet / lzy4 pgm* young lateral root without gravistimulation. Time-lapse microscopy was performed on the vertical stage confocal microscope. Images were acquired at 20-second intervals for 15 min. *g*, the direction of gravity.

**Movie S4.**

Columella cells of *LZY4p::LZY4-mScarlet / lzy4* primary root with gravistimulation. Time-lapse microscopy was performed on the vertical stage confocal microscope with seedling grown in the micro hydroponic chamber. Images were acquired at 20-second intervals for 15 min. *g*, the direction of gravity. Image acquisition was initiated immediately after gravistimulation by rotating the stage 135 degrees.

**Movie S5.**

Columella cells of *LZY4p::LZY4-mEos2 / lzy4* young lateral root. Time-lapse microscopy was performed on the vertical stage confocal microscope. Images were acquired at 20-second intervals for 15 min. *g*, the direction of gravity. Image acquisition was initiated immediately after photo-conversion by irradiation of a 405 nm laser.

**Movie S6.**

Irradiation with a laser for optical tweezer at around the plasma membrane of the columella cell without manipulating amyloplasts (fig. S15, D and E). Z stack images of LZY4-mScarlet fluorescence were acquired before and after the laser irradiation, and max intensity projection images were then produced.

**Movie S7.**

Amyloplast manipulation from the lower to the upper side of the columella cell with the optical tweezer (fig. S15, J and K). Z stack images of LZY4-mScarlet fluorescence were acquired before and after the manipulation, and max intensity projection images were then produced.

**Movie S8.**

Changes in LZY4-mScarlet fluorescence intensity on the plasma membrane in response to amyloplast manipulation with the optical tweezer (Fig. 4C). LZY4-mScarlet fluorescence (left) and bright-field images (right) are shown.

**Table S1. (separate file)**

List of primers used in this study.
